# Layer-specific inhibitory microcircuits of layer 6 interneurons in rat prefrontal cortex

**DOI:** 10.1101/2020.01.27.920686

**Authors:** Chao Ding, Vishalini Emmenegger, Kim Schaffrath, Dirk Feldmeyer

## Abstract

GABAergic interneurons in different cortical areas play important roles in diverse higher order cognitive functions. The heterogeneity of interneurons is well characterized in different sensory cortices, in particular in primary somatosensory and visual cortex. However, the structural and functional properties of the medial prefrontal cortex (mPFC) interneurons have received less attention. In this study, a cluster analysis based on axonal projection patterns revealed four distinct clusters of L6 interneurons in rat mPFC: Cluster 1 interneurons showed axonal projections similar to Martinotti-like cells extending to layer 1, cluster 2 displayed translaminar projections mostly to layer 5, cluster 3 interneuron axons were confined to the layer 6, whereas those of cluster 4 interneurons extend also into the white matter. Correlations were found between neuron location and axonal distribution in all clusters. Moreover, all cluster 1 L6 interneurons showed a monotonically adapting firing pattern with an initial high-frequency burst. All cluster 2 interneurons were fast-spiking, while neurons in cluster 3 and 4 showed heterogeneous firing patterns. Our data suggest that L6 interneurons that have distinct morphological and physiological characteristics are likely to innervate different targets in mPFC thus play differential roles in the L6 microcircuitry and in mPFC-associated functions.

## Introduction

Seminal research in rats, monkeys and humans has revealed that the prefrontal cortex (PFC) plays a significant role in executive function including cognitive processes such as memory, attention, and decision-making (Fuster 2001; Miller and Cohen 2001; Euston et al. 2012). While the rodent PFC is undoubtedly less complex than its human counterpart, it nevertheless encompasses similar executive functions. The rodent PFC acts like a nodal station of cortical networks by receiving multimodal cortico-cortical projections, such as from motor, somatosensory, gustatory, limbic and auditory cortices (Uylings et al. 2003a). The medial prefrontal cortex (mPFC), which includes the anterior cingulate cortex, prelimbic cortex and infralimbic cortex, consists of 80-90% glutamatergic pyramidal cells and 10-20% GABAergic interneurons (Riga et al. 2014). Analogous to interneurons in sensory cortices, mPFC interneurons can be distinguished based on their morphological, functional, and molecular properties (for reviews s. Markram et al. 2004; Ascoli et al. 2008; Defelipe et al. 2013; Kepecs and Fishell 2014; Lein et al. 2017; Zeng and Sanes 2017; Yuste et al. 2019).

GABAergic interneurons play a significant role in orchestrating network activity by regulating the concerted activity of pyramidal cells (for reviews see Tremblay et al. 2016). Different types of interneurons innervate different subcellular compartments of pyramidal cells, thereby regulating their activity (Palmer et al. 2012). The intricate balance between excitatory and inhibitory neurotransmission (the so-called E-I balance) plays a fundamental role in executing a range of PFC-dependent behaviours (Ferguson and Gao 2018). Disturbances in the inhibitory regulation of cortical activity can have dramatic consequences resulting in neuropsychiatric conditions, amongst them schizophrenia, anxiety, or depression (Benes and Berretta 2001; Volk and Lewis 2005; Beneyto et al. 2011; Möhler 2012).

In order to understand neuronal circuit dynamics, it is crucial to investigate the different neuronal cell types and their connectivity pattern in different layers. While the heterogeneity of pyramidal cells and interneurons in sensory and motor cortices is well characterized (Yang et al. 1996; Markram et al. 2004; Helmstaedter et al. 2009; Rudy et al. 2011; Defelipe et al. 2013; Marx and Feldmeyer 2013; Tremblay et al. 2016; Emmenegger et al. 2018; Feldmeyer et al. 2018), only few studies are currently available investigating the diversity of interneurons in mPFC (Ahrlund-Richter et al. 2019; Sun et al. 2019). The deepest layer of mPFC received less attention in classifying its interneuron complement. This is surprising as the layer 6 (L6) of mPFC emerges as a key player in attention control (Wimmer et al. 2015). L6 excitatory neurons project extensively to thalamic nuclei and to local circuit interneurons in cerebral cortex (Guillery and Sherman 2002; Thomson et al. 2002; West et al. 2006; Zikopoulos and Barbas 2006; Thomson 2010; Arzt et al. 2018). Yet, the axonal projection motifs and connectivity profiles of L6 interneurons in other cortical areas have received less attention. To understand the complexity of the L6 microcircuitry, it is pivotal to unravel the interneuron complement of this layer. Quantitative measures of axon projections are considered to be of high functional significance because they are one of the parameters defining neuronal innervation domains thereby making a reliable prediction of synaptic connectivity possible (Lübke et al. 2003; Feldmeyer et al. 2018). Therefore, we used axonal projections with reference to layer borders and soma position as primary classifiers and employed objective classification methods such as unsupervised cluster analysis to identify different interneuron types in L6 of rat mPFC. We found that L6 interneurons display layer-specific axonal projections, for instance local L6 projecting interneurons and translaminar projecting interneurons, which were further subdivided into locally projecting interneurons, L6/white-matter (WM) projecting interneurons, L5 projecting interneurons, L1/2/3 projecting interneurons. Correlation between electrophysiological properties and axonal projection pattern was found in some of the interneuron types. We suggest that L6 interneurons in mPFC with distinct morphologies innervate pyramidal cells in different layers and even different subcellular compartments and may therefore play different roles in modulating excitatory signalling. In this way they may contribute to the maintenance of the excitation-inhibition (E-I) balance in the prefrontal and associated cortical networks.

## Methods

### Slice preparation

All experimental procedures were carried out in accordance with the guidelines of the Federation of European Laboratory Animal Science Association (FELASA), the EU Directive 2010/63/EU, and the German animal welfare act, and were approved by the Northrhine-Westphalian Landesamt für Natur-, und Verbraucherschutz (LANUV).

In this study, Wistar rats (Charles River, either sex) aged 17-21 postnatal days (P17-21) were used. The experimental procedures have been described previously (Van Aerde and Feldmeyer 2015) with minor modifications. Briefly, rats were deeply anesthetized with isoflurane and decapitated. The brain was quickly removed and placed in an ice-cold artificial cerebrospinal fluid (ACSF) containing: 125 mM NaCl, 2.5 mM KCl, 1.25 mM NaH_2_PO_4_, 4 mM MgCl_2_, 1 mM CaCl_2_, 25 mM NaHCO_3_, 25 mM glucose, 3 mM Myo-Inositol, 2 mM Na-pyruvate, and 0.4 mM ascorbic acid; the osmolarity of the solution was ∼310 mOsm. The concentration of Ca^2+^ was lowered to reduce potential excitotoxic synaptic transmission during slicing. In order to maintain adequate oxygenation and a physiological pH level, the solution was constantly bubbled with carbogen gas (95% O_2_ and 5% CO_2_). Coronal slices of the prelimbic mPFC at 350 µm thickness were made using a vibrating microslicer at a low speed and high vibration frequencies. The slices were then transferred to an incubation chamber for a recovery period of ∼1 hr at room temperature.

During whole-cell patch-clamp recordings, slices were continuously perfused (perfusion speed ∼5 ml/min) with an artificial ACSF containing: 125 mM NaCl, 2.5 mM KCl, 1.25 mM NaH_2_PO_4_, 1 mM MgCl_2_, 2 mM CaCl_2_, 25 mM NaHCO_3_, and 25 mM glucose, bubbled with carbogen gas and maintained at ∼31°C. Patch pipettes were pulled from thick-wall borosilicate glass capillaries and filled with an internal solution containing: 135 mM K-gluconate, 4 mM KCl, 10 mM HEPES, 10 mM phosphocreatine, 4 mM Mg-ATP, and 0.3 mM GTP (pH 7.4 with KOH, osmolarity ∼300 mOsm). In order to obtain permanent stainings of the patched neurons for the morphological analysis, 13.4 mM biocytin was added to the internal solution.

Neurons were visualized using infrared differential interference contrast microscopy. The layers were distinguished based on the cell density and cell soma size in agreement with earlier studies on the prefrontal cortex (Van Eden and Uylings 1985; Gabbott et al. 1997; Uylings et al. 2003b; Gabbott et al. 2005; Van Aerde and Feldmeyer 2015). Overall, the prefrontal cortex can be divided into three sections - the upper third comprises L1-L3, the middle third includes L5 and the lower third constitutes L6. Unlike other cortical regions, L1 in prefrontal cortex is distinctly wide. L2 is the thinnest layer of the prefrontal cortex, almost devoid of neurons except few interneuron types, therefore seen as a thin dark band between L1 and L3. Despite the lack of a granular layer, L3 and L5 is demarcated by a band of thalamocortical axon collaterals in deep L3 (Uylings and Van Eden 1990; Uylings et al. 2003b; Kubota et al. 2007; Hirai et al. 2012)

### Electrophysiology

Whole-cell patch clamp recordings were made using an EPC10 amplifier (HEKA, Lambrecht, Germany). Signals were sampled at 10 kHz, filtered at 2.9 kHz using Patchmaster software (HEKA), and later analyzed off-line using Igor Pro software (Wavemetrics, USA). The recordings were performed using patch pipettes of resistance between 5 to 7 MΩ. After establishing the whole-cell configuration, the resting membrane potential was measured immediately after breakthrough. Bridge balance and capacitance neutralization were adjusted. Whole-cell series resistance was monitored throughout the experiment and was compensated by 80%. Recordings with a series resistance exceeding 50 MΩ were excluded from the data analysis. Membrane potentials were not corrected for the junction potential. Passive and active action potential properties were assessed by eliciting a series of initial hyperpolarizing, followed by depolarizing current pulses under current clamp configuration for 1 s. We used the intrinsic firing properties recorded in the current clamp mode to differentiate interneurons from pyramidal cells, and morphological reconstructions for a post-hoc confirmation.

### Histological Procedures

After the electrophysiological recordings, slices containing biocytin-filled neurons were fixed at 4 °C in 100 mM phosphate buffer solution (PBS, pH 7.4) containing 4% paraformaldehyde (PFA) for at least 24 h. After rinsing slice several times in PBS, slices were treated with 1% H_2_O_2_ in PBS for about 20 min, in order to reduce any endogenous peroxidase activity. Following this slices were rinsed repeatedly using 100 mM PBS, then incubated in 1% avidin-biotinylated horseradish peroxidase (Vector ABC staining kit, Vector Lab. Inc.) containing 0.1% Triton X-100 for 1 h at room temperature. This was followed by a chromogenic reaction that resulted in a dark precipitate by adding 0.5 mg/ml 3,3-diaminobenzidine (DAB; Sigma-Aldrich, USA) until distinct axonal and dendritic branches of the biocytin-filled neurons were clearly visible. The slices were rinsed again with 100 mM PBS, followed by slow dehydration in increasing ethanol concentrations and finally in xylene for 2-4 h (Marx et al. 2012). Slices were then mounted on gelatinized slides and embedded using Eukitt medium (Otto Kindler GmbH).

### Immunohistochemical staining

For the identification of molecular markers expressed by GABAergic interneurons, 100 µm thin brain slices were prepared using a vibratome. Brain slices were immediately fixed with 4% PFA in 100 mM PBS for ∼4 h at 4 °C. The slices were then permeabilised in 0.5% Triton X-100 in 100 mM PBS with 1% milk powder for 1 h at room temperature. Primary and secondary antibodies for parvalbumin (PV), somatostatin (SOM) and the transcription factor Prox1 were diluted in a permeabilisation solution containing 0.5% Triton X-100 and 100 mM PBS. Subsequently, slices were incubated overnight in a solution containing the respective primary antibody at 4 °C and then rinsed thoroughly with 100 mM PBS. Slices were then treated with the secondary antibodies for 2-3 h at room temperature in the dark. Controls were also performed in the absence of primary and secondary antibodies. After being rinsed in 100 mM PBS for several times, slices were embedded in Moviol and visualised by transmitted light fluorescence microscopy using an upright microscope equipped with fluorescence optics. The fluorescence images were taken using the Olympus CellSens platform.

In a subset of experiments, we tried to identify the expression of molecular markers in single L6 interneurons in brain slices to investigate a possible correlation with the immunohistochemical, morphological and electrophysiological properties. To this end, Alexa Fluor® 594 biocytin salt (1:500, Invitrogen, Darmstadt, Germany) was added to the internal solution (composition as described above) to identify the patched neurons in the post hoc antibody labelling methods. After recording, slices were fixed in 4% PFA in 100 mM PBS for 2 to 4 h at 4 °C and antibody labelling was performed as described above. The location of the stained neuron in the slice was visualized by the conjugated Alexa dye, so that the expression of a specific molecular marker could be tested in identified individual neurons. After acquiring fluorescent images, slices were incubated in 100 mM PBS overnight and subsequently processed for morphological analyses (see above).

### Morphological 3D Reconstructions

Morphological reconstructions of biocytin filled Layer 6 mPFC interneurons were made using Neurolucida^®^ software (MicroBrightField, Williston, VT, USA) on an upright microscope equipped with a motorized stage at a magnification of 100-fold using oil-immersion objective. Neurons were selected for reconstruction based on the quality of biocytin labelling, in particular when background staining was minimal. Embedding using Eukitt medium reduced fading of cytoarchitectonic features and enhanced contrast between layers (Marx et al. 2012). This allowed the reconstruction of different layer borders along with the neuronal reconstructions. Furthermore, the position of soma and layers were confirmed by superimposing the DIC images taken during the recording. All reconstructions were aligned with respect to the pial surface. The tissue shrinkage was corrected for using shrinkage correction factors of 1.1 in the *x*–*y* direction and 2.1 in the *z* direction (Marx et al. 2012).

### Data Analysis

#### Membrane and Spike Properties

Custom written macros for Igor Pro 6 (Wavemetrics) were used for the analysis of the recorded electrophysiological data.

The resting membrane potential (V_rest_) of the neuron was measured directly after breakthrough into the whole-cell configuration with no current injection. To calculate the input resistance (R_in_), the slope of the linear fit to the voltage step from −60 mV to −70 mV of the current-voltage (I-V) relationship was used. Rheobase current was defined as the minimal current that elicited the first spike.

For the analysis of a single spike characteristics such as threshold, amplitude and half-width, a smaller step size increment (10 pA) for current injection was used to ensure that the action potential elicited is very close to its potential threshold. The spike threshold was defined as the point of maximal acceleration of the membrane potential using the second derivative (d_2_V/dt^2^), i.e. the time point with the fastest voltage change. The spike amplitude was calculated as the difference in voltage from AP threshold to the peak during depolarization. The spike half-width was measured as the time difference between rising phase and decaying phase of the spike at half-maximum amplitude. Inter-spike interval (ISI) was measured as the average time taken between individual spikes at the current step that elicited close to 10 APs. The adaptation ratio was measured as the ratio of the tenth ISI and the second ISI.

#### Morphological Properties

In order to ensure high-quality reconstructions, the degree of truncations were carefully examined. If the primary branch of an axon showed immediate truncation, or the total length of axons were less than 7 mm, neurons were excluded from the data analysis.

3D reconstructed neurons were quantitatively analysed using Neuroexplorer® software (MicroBrightField, Williston, VT, USA), Morphological properties such as the distributions of axonal length in cortical layer 1 to 6, the white matter, and the axonal orientation were extracted from Neuroexplorer® software. The axonal orientation was calculated using polar histogram, where the spatial geometry is transformed into length and direction and the ratio of the axon length in the horizontal and vertical directions was subsequently calculated determined.

#### Unsupervised Hierarchical Cluster Analysis

In order to perform an objective classification of interneuron subtypes, we used unsupervised hierarchical cluster analysis using Ward’s method (Ward 1963). This method utilizes a minimum variance method to combine cells into clusters at each stage, which minimizes the total within-cluster variance. Euclidean distance was used to calculate the variance. A dendrogram was constructed to visualize the distance at which clusters are combined.

#### 2D Axon Distribution Maps and Vertical Axon Distribution Profiles

2D and 1D axon distribution maps were made by using custom-written software in Matlab (courtesy of Drs. G. Qi and H. Wang). 3D maps of axonal density were obtained using computerised 3D reconstructions. The soma centre of each neuron in a single cluster was aligned and given the coordinates of X, Y, Z = (0, 0, 0). By using the segment point analysis in the Neuroexplorer^®^, the relative coordinate of the beginning and endpoint of each segment in the axonal trace were acquired. The 3D density map for a cluster was constructed for each reconstructed neurons in this cluster and then averaged in Matlab. The averaged density maps were smoothed using the 3D smoothing function in Matlab with a Gaussian kernel (s.d = 50 µm) and isosurfaces were calculated at the 80 percentile. 2D axon distribution maps were made by showing the 3D map in a 2D way with only x axis and y axis. 1D axon density profiles were calculated at a resolution of 50 µm by extracting value of y axis. The average curve of single group was made by aligning the soma position of individual profile.

### Statistics

All data are represented as mean ± standard deviation. Because the distribution of measured parameters was not normal, the *Kruskal-Wallis test* was used for statistical comparisons among all clusters; for comparison between two independent groups, *Wilcoxon-Mann-Whitney test* was used. Correlation analysis was performed by calculating Pearson correlation coefficients.

## Results

L6 interneurons in rat prefrontal cortex were recorded using whole-cell patch-clamp with simultaneous biocytin filling. After recording and histochemistry, neurons were selected for further analysis if no apical dendrite was visible. A low spine density was used as an additional criterion to distinguish interneurons from excitatory neurons. In this study, the average spine density per 10 µm dendrite was found to be 0.29 ± 0.18 for all types of interneurons, a value significantly lower than that for pyramidal cells (Elston 2003; Arzt et al. 2018). In a subset of experiments in which the expression of specific interneuron marker proteins was tested, the red fluorescent dye Alexa 594 was added together with biocytin to identify the location of the patched neurons prior to antibody labelling.

### Morphological Classification of L6 interneurons

All L6 interneurons with an axon length exceeding 7 mm were selected for further analysis, resulting in a total of 48 high-quality 3D morphological reconstructions. Quantitative morphological classification of L6 interneurons was performed based on their axonal projection patterns. Specifically, parameters used for cluster analysis were the distribution of the axonal length in cortical layers 1 to 6 and the white matter (WM), the axon orientation, and the axonal distribution with respect to the soma position in layer 6 (from the pial surface to the soma and from the soma to the WM; see Methods). Based on these morphological parameters, four clusters were identified with distinct axonal projection patterns: L1/2/3-projecting interneurons, L5-projecting interneurons, locally-projecting L6 interneurons and L6/WM-projecting interneurons (Fig. 1A, Suppl. Fig. 1 and 2).

**Figure 1.**
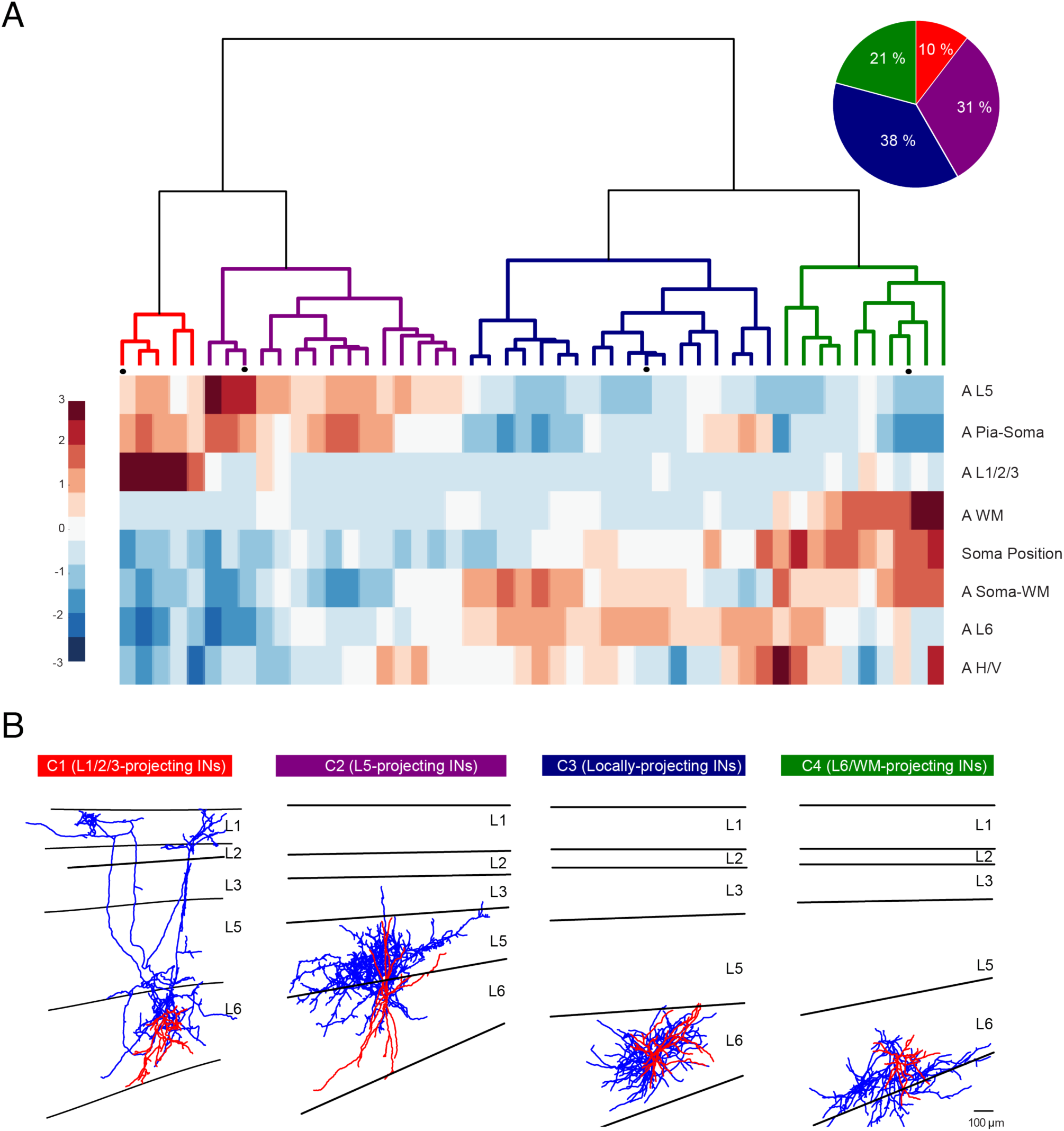
Morphological analysis of L6 inhibitory neurons using unsupervised cluster analysis. (A) Dendrogram obtained from a cluster analysis based on morphological parameters reveals four clusters of L6 interneurons (n=48). The color map below the dendrogram indicates the standardized values of the corresponding parameters (listed on the right) of individual neurons, in which red represents values above the mean, white represents the mean, and blue represents values below the mean. Pie chart shows the percentage contribution of each cluster. (B) Representative axodendritic morphologies of the four clusters. Dendrites are shown in red and axons in blue. Scale bar is 100 µm. A L5: Axonal distribution in L5; A Pia-Soma: Axonal distribution from Pia-Soma; A L1/2/3: Axonal distribution in L1-L3; A WM: Axonal distribution in WM; A Soma-WM: Axonal distribution from Soma-WM; Soma Position: Relative soma position in L6; A L6: Axonal distribution in L6; A H/V: Axon horizontal/vertical).

### Morphological Cluster 1: L1/2/3-projecting interneurons

Of the 48 interneurons, five were L1/2/3-projecting interneurons constituting 10% of the population. All showed a largely vertical axonal projection pattern (axon horizontal/vertical: 0.73 ± 0.16, example in Fig. 1B, all reconstructions in Suppl. Fig. 2A). Compared to PFC L6 interneurons in clusters 2-4 (see below), a significantly larger fraction of the axonal collaterals of cluster 1 L6 interneurons resided in L1 to L3 (37.2 ± 7.5%, P = 5.9E-04 between all clusters; Table 1). However, the axon plexus in L6 was significantly smaller (20.8 ± 16.2%, P = 1.16E-12 between all clusters; Table 1). All cluster 1 L6 interneurons had axon collaterals terminating in L1. Two of them showed horizontally projecting collaterals in L1, while the other three cluster 1 interneurons sent several collaterals to L5 and L3. The relative soma position of these PFC L6 interneurons in their home layer was calculated as the ratio between the distance from soma to the L5/L6 border and the vertical length of L6. L1/2/3- projecting interneurons had an average soma position value of 0.15 ± 0.09, i.e. they were located near the L5/L6 border; their axon collaterals projected mostly towards the pial surface (ratio of axonal distribution from pia to soma: 0.87 ± 0.09, Fig 2A and B).

**Figure 2.**
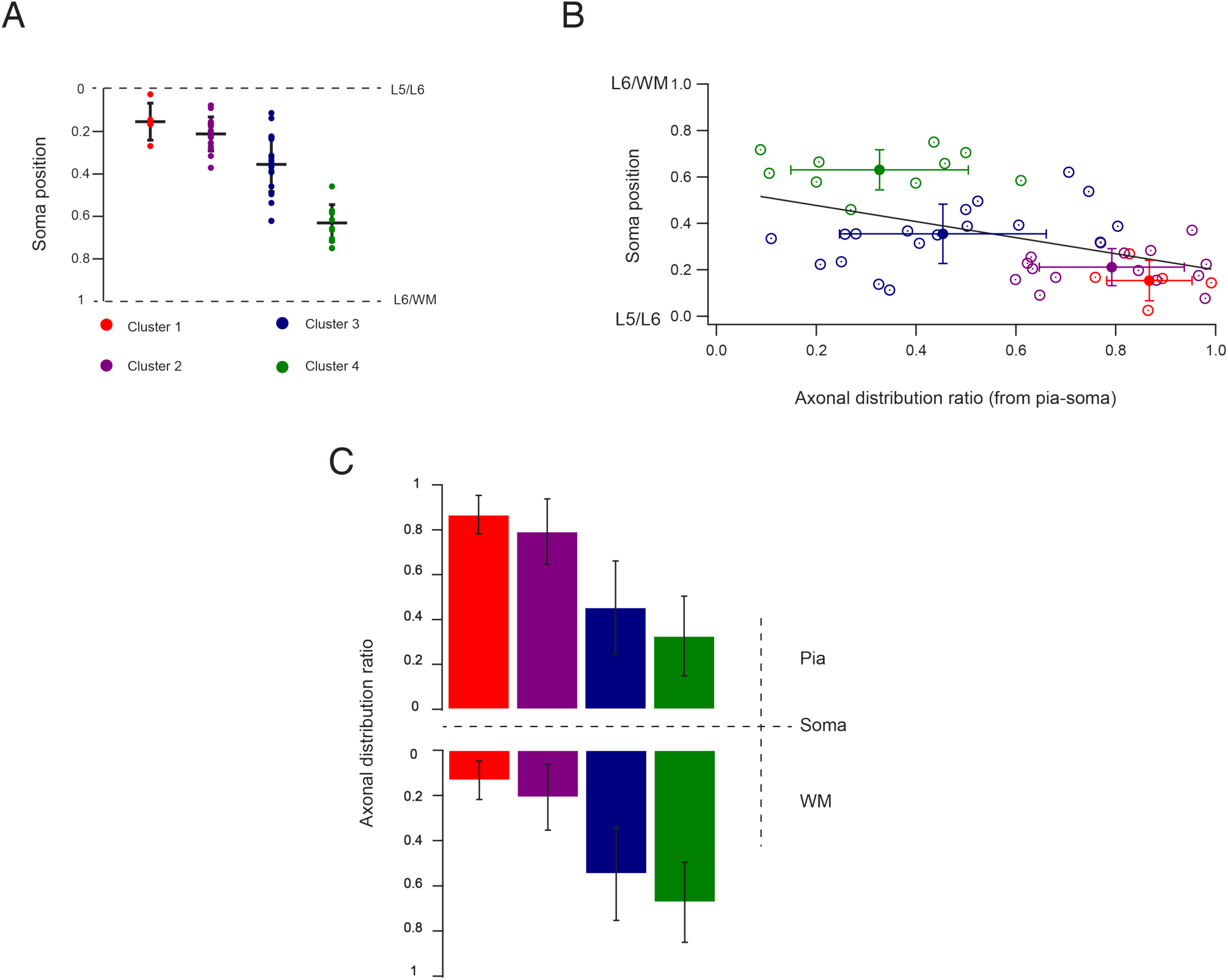
mPFC L6 interneurons display a strong correlation between soma position and axonal projection pattern. (A) Relative soma position in L6 of interneurons of the four morphological clusters. (B) Correlation between soma position of individual L6 interneurons and their axonal distribution from pia to soma. (C) Relative axonal distributions of the neurons from different clusters are shown. Top: from pia to soma; Bottom: from soma to WM.

**Table 1.**
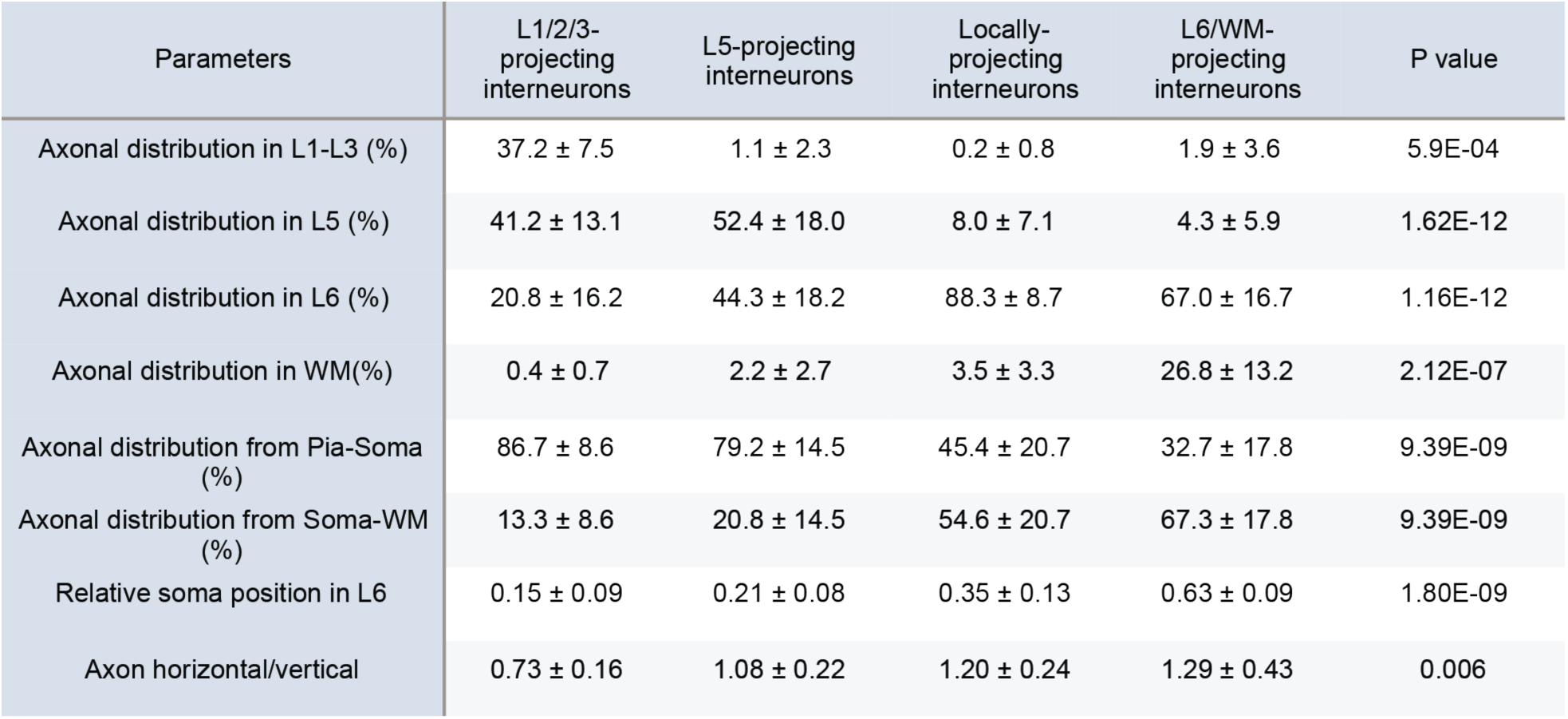
Statistical analysis of the axonal properties of L6 inhibitory neurons in the four morphological clusters. All data are represented as mean ± standard deviation. The distribution of the axon in the different layers are given as percentage values. The Kruskal-Wallis test was used to test for the significant difference between clusters.

### Morphological Cluster 2: L5-projecting interneurons

The second cluster, the L5-projecting interneurons, consisted of 15 interneurons and constituted 31% of the L6 interneuron population. Like L1/2/3-projecting interneurons, these interneurons were also located near the L5/6 border (relative soma position: 0.21 ± 0.08, Fig 2A). Of all axon collaterals 79.2 ± 14.5% projected towards the pial surface (Fig 2, B), mostly terminating in L5 with a dense axonal pattern (example in Fig. 1, B, density map in Fig. 3, all reconstructions in Suppl. Fig. 1B). L5-projecting interneurons preferentially innervated L5 of PFC: of all PFC L6 interneuron clusters, these neurons had the largest fraction of the axon residing in L5 (52.4 ± 18.0%, P = 1.62E-12 between all clusters, Table 1).

**Figure 3.**
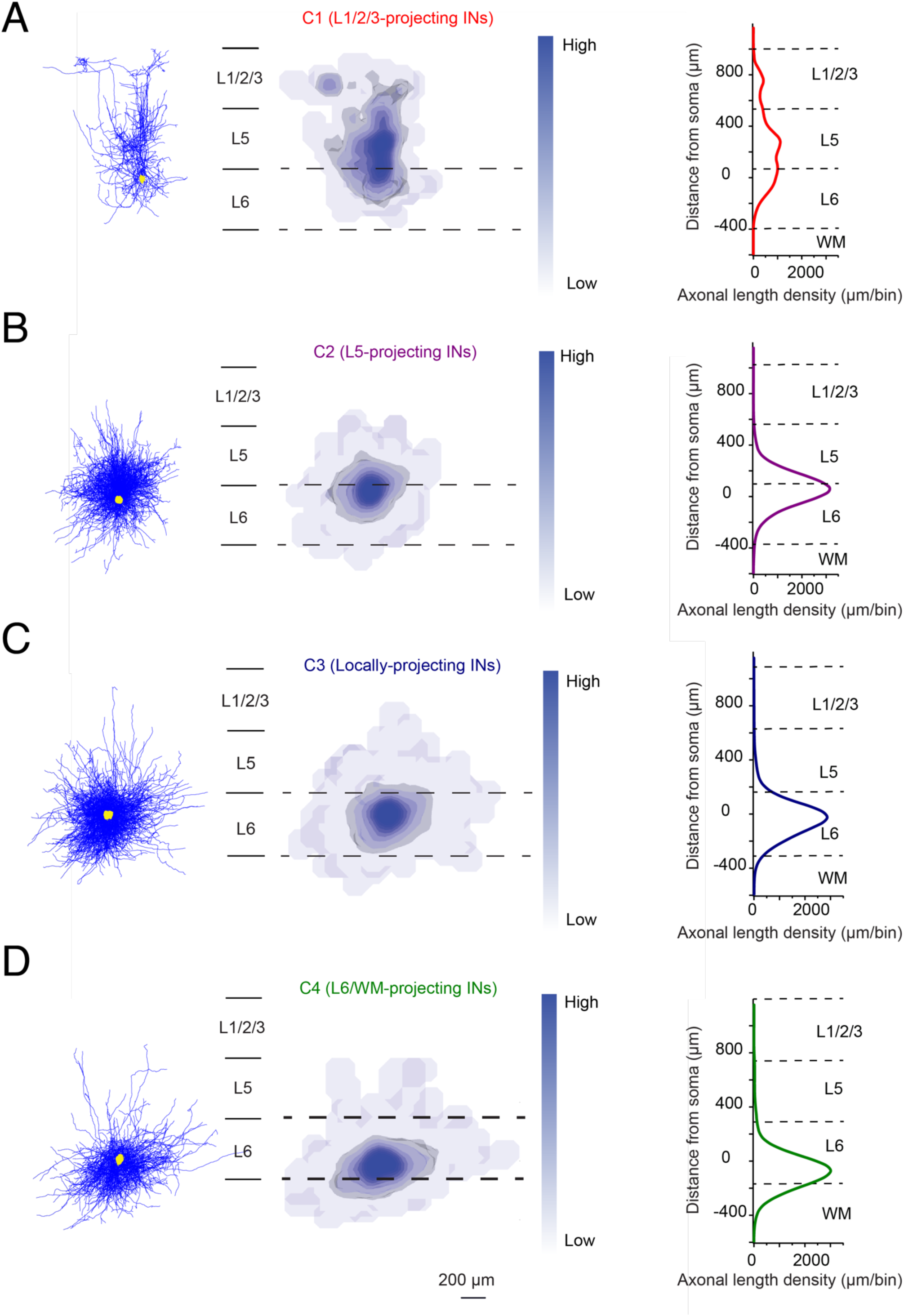
Reconstruction overlays and density maps of inhibitory mPFC L6 interneuron axons based on 4 morphological clusters. Left: All interneurons are aligned with their soma position represented as yellow dots; axons are shown in blue. Scale bar = 200 µm. Middle: 2D density maps of different clusters, the gray isosurfaces show the 80 percentile of the axonal density. Right: vertical axonal distribution of L6 interneuron clusters. The individual curves show the average axon density distribution along the vertical axis; bin size in the x axis: 50 µm in vertical direction. Dashed lines indicate layer borders. Results from 5 L1/2/3-projecting interneurons, 15 L5-projecting interneurons, 18 locally-projecting L6 interneurons and 10 L6/WM-projecting interneurons are shown in panel A, B, C and D respectively.

### Morphological Cluster 3: Locally-projecting interneurons

Cluster 3 of PFC L6 interneurons, termed locally-projecting L6 interneurons consisted of 18 neurons, i.e. 38% of the total number of interneurons (all reconstructions in Suppl. Fig. 2A). Axons of these interneurons were confined to their home layer with a distribution ratio of 88.3 ± 8.7%, a value significantly larger than that of all other clusters (P = 1.16E-12 between all clusters, Table 1 and Fig.3). Locally-projecting L6 interneurons were found in the middle of L6 with an average soma position of 0.35 ± 0.13; their axons showed no preferential projection towards superficial or deep cortical layers (45.4 ± 20.7% of the axons found between pia and soma while 54.6 ± 20.7% were located between soma-and WM, Fig 2A and B). This projection pattern suggests that locally-projecting L6 interneurons mainly innervate a narrow stratum in their home layer.

### Morphological Cluster 4: L6/WM-projecting interneurons

The last cluster, the L6/WM-projecting interneurons, comprised 10 cells and constituted 21% of the total number of interneurons in our study (all reconstructions in Suppl. Fig. 2B). Interneurons of this cluster displayed a dominant horizontal axonal projection compared to all other clusters (axon horizontal/vertical: 1.29 ± 0.43, P = 0.006 between all clusters, Table 1). They were found closer to the WM rather than the L5/L6 border (relative soma position: 0.63 ± 0.09, P = 1.8E-09 compared to all other clusters, Table 1 and Fig. 2A); therefore, they are mostly L6B interneurons: The axon collaterals of these neuron cluster projected preferentially towards the WM and less so to the pial surface (67.3 ± 17.8 vs. 32.7 ± 17.8, Fig 2B). Unlike all other clusters, a significant proportion of the axon of cluster 4 L6 interneurons was located in the WM (26.8 ± 13.2%, P = 2.12E-07 when compared to all the other clusters, Table 1 and Fig. 3).

The axonal projection pattern of L6 interneurons in PFC was correlated with the position of the interneuron soma in L6 (P = 0.0005). Interneurons located at the L5/L6 border have axons that project preferentially to the pial surface with only few collaterals projecting to the WM (Fig. 2C). In contrast, neurons located close to the L6/WM border extended their axons predominately in the direction of the WM (Fig. 2D).

### Correlation between morphological and electrophysiological properties

Several studies have demonstrated that the axonal projection pattern of cortical interneurons and their intrinsic electrophysiological properties are - if at all - only weakly correlated so that a combination of electrophysiological and morphological parameters is often not helpful to classify interneurons (Gupta et al. 2000; Wang et al. 2002; Markram et al. 2004; Wang et al. 2004; Kumar and Ohana 2008; Helmstaedter et al. 2009; Arzt et al. 2018; Emmenegger et al. 2018; Feldmeyer et al. 2018). In this study, active electrical properties of interneurons in the morphological clusters were determined and compared for different morphological clusters. Interneurons with a series resistance of >50 MΩ were excluded from this analysis. Therefore, in order to determine a correlation between electrophysiological and morphological parameters we used 5 neurons in morphological C1, 10 in C2, 11 in C3 and 9 in C4. PFC L6 interneurons could be categorized into several groups based on their firing behavior (Ascoli et al. 2008; Feldmeyer et al. 2018). Here, we found that L1/2/3-projecting interneurons were adapting neurons (example in Fig. 4A and B) with a significantly smaller adaptation ratio of 0.38 ± 0.24 (2_nd_ ISI / 10_th_ ISI; P = 7.08E-04, Table 2, details of comparison in Fig. 4C) and a larger standard deviation of the ISI (34.1 ± 13.7 ms; P = 2.18E-04, Table 2, details of comparison in Fig. 4C) compared to interneurons of all other clusters. Moreover, the AP halfwidth of L1/2/3-projecting interneurons was much longer compared to interneurons in morphological clusters 2 to 4 (0.81 ± 0.16 ms; P = 0.001, Table 2, details of comparison in Fig. 4C). All L5-projecting interneurons displayed a fast-spiking (FS) firing pattern; eight of them displayed stuttering firing behavior, while the other seven showed continuous AP firing. (examples in Fig. 4A and B). The ISI of L5-projecting interneurons was 26.0 ± 10.2 ms, which was the smallest among all four clusters (P = 0.008, Table 2, details of comparison in Fig. 4C). In addition, these L6 interneurons had also the largest AHP amplitude (25.2 ± 3.4 mV; P = 0.001, Table 2, details of comparison in Fig. 4C). Unlike interneurons in clusters 1 and 2, those in clusters 3 and 4 displayed heterogeneous firing patterns (but see below). Interneurons with FS, adapting and non-adapting non-FS firing behavior were found in these two clusters (examples in Fig. 4A and B).

**Figure 4.**
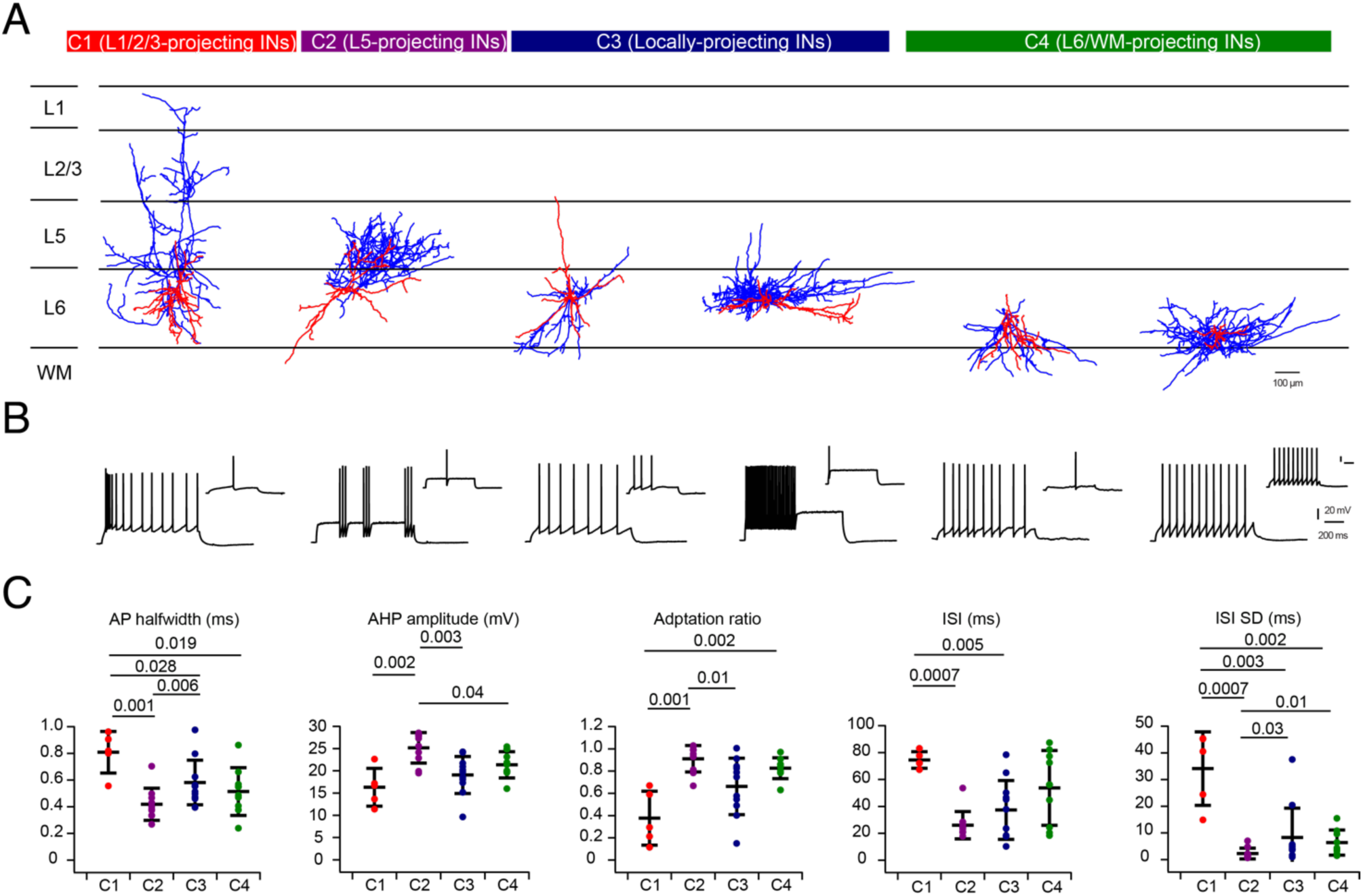
Comparison of electrophysiological parameters in morphological clusters of L6 interneurons in rat mPFC. (A) Representative examples of L6 interneurons of the four morphological clusters. Axons are given in blue, the somatodendritic domain in red. (B) Corresponding firing pattern of the neurons that are shown in A (10-spike train and the first AP at rheobase current injection). (C) Statistical analysis of the active electrophysiological properties of mPFC L6 interneurons in the four morphological clusters. A Wilcoxon-Mann-Whitney test was performed for the significant difference among clusters. P value are shown above.

**Table 2.** Statistical analysis of the electrophysiological parameters under 4 morphological clusters. All data are represented as mean ± standard deviation. The Kruskal-Wallis test was used to test for significant differences between clusters.

| Parameters | L1/2/3-projecting interneurons | L5-projecting interneurons | Locally-projecting interneurons | L6/WM-projecting interneurons | P value |
| --- | --- | --- | --- | --- | --- |
| ISI average (ms) | 74.5 ± 6.1 | 26.0 ± 10.2 | 37.3 ± 21.9 | 53.7 ± 27.8 | 0.008 |
| ISI SD (ms) | 34.1 ± 13.7 | 2.3 ± 2.0 | 8.3 ± 11.0 | 6.4 ± 4.7 | 2.18E-04 |
| AP half-width (ms) | 0.81 ± 0.16 | 0.42 ± 0.12 | 0.58 ± 0.17 | 0.51 ± 0.18 | 1.45E-03 |
| AP threshold (mV) | -37.9 ± 1.9 | -30.6 ± 6.2 | -32.2 ± 7.5 | -33.7 ± 3.3 | 0.04 |
| AHP amplitude (mV) | 16.3 ± 4.2 | 25.2 ± 3.4 | 19.1 ± 4.2 | 21.4 ± 2.9 | 1.22E-03 |
| Adaptation ratio | 0.38 ± 0.24 | 0.91 ± 0.12 | 0.66 ± 0.25 | 0.83 ± 0.09 | 7.08E-04 |
| Firing Frequency (Hz) / 100 pA | 24.1 ± 6.3 | 36.7 ± 14.9 | 30.8 ± 21.1 | 32.5 ± 14.4 | 0.39 |

To identify a possible correlation between axonal projecting pattern and AP firing properties, we performed a CA with the 35 interneurons with both morphological and electrophysiological properties that matched the criteria given above. The morphological and electrophysiological parameters used for the CA are listed in Fig. 5A. The combined CA revealed four clusters, - termed clusters A-D - which were largely similar to the morphological cluster 1-4 (Fig. 5A and Fig. 1A). Cluster A consisted of 5 L1/2/3-projecting interneurons and one locally-projecting (cluster 3) L6 interneurons. Cluster B comprised 11 interneurons, including all the L5-projecting interneurons and one locally-projecting interneurons. Cluster C consisted of 9 locally-projecting interneurons and one L6/WM-projecting interneurons, while the remainder of the L6/WM-projecting interneurons were placed in cluster D. A strong correlation between axonal morphology and active membrane properties was found for cluster A and B; for cluster C and D sub-clusters showing either a FS or a non-FS firing patterns were identified (Suppl. Fig. 3).

**Figure 5.**
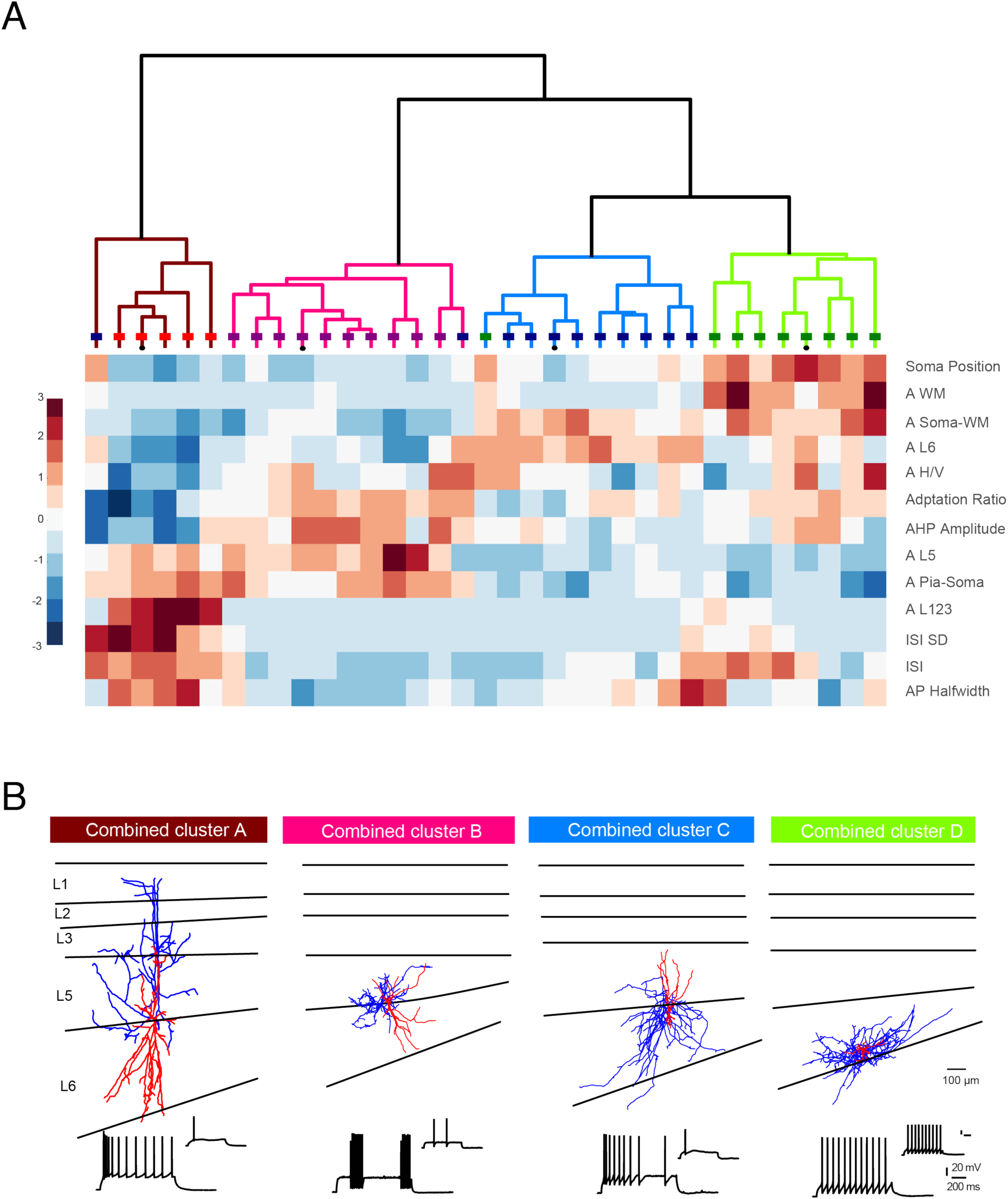
Comparison of morphological and electrophysiological parameters of L6 interneurons in rat mPFC. (A) A combined unsupervised hierarchical cluster analysis based on both morphological and electrophysiological parameters revealed four clusters. The cut-off for significant clusters is 70% of the maximum linkage distance. Colored lines in each dendrogram represent the color code from the morphological clusters in (B). Colored boxes on each line shows the morphology clusters the neurons belong to. The color map below the dendrogram indicates the standardized values of the corresponding parameters (listed on the right) of individual neurons, in which red represents values above the mean, white represents the mean, and blue represents values below the mean. (see Fig. 1). (B) Representative examples of L6 inhibitory neurons based on combined morphological/electrophysiological parameters. Axons are given in blue, the somatodendritic domains in red. Corresponding firing pattern are shown below each example morphology.

#### Neurochemical expression of interneurons in L6 of mPFC

To correlate the electrophysiological properties with the neurochemical maker expression specific for different cortical interneuron types, we performed patch-clamp recordings with simultaneous filling of biocytin and the fluorescent dye Alexa 594 to allow an unambiguous identification of the recorded neuron. After determining the passive and active electrophysiological properties, slices were briefly fixed in PFA and processed for immunostaining for PV, SOM and Prox1, a marker for vasoactive intestinal peptide (VIP)-expressing interneurons and a fraction of reelin-and calretinin-expressing interneurons (Rubin and Kessaris, 2013; Miyoshi et al., 2015). A subsequent DAB staining of the slices was made to determine the morphology of labeled neurons. Of 17 PFC L6 interneurons, seven were PV-positive, four SOM-positive and five Prox1-positive. One interneuron showed no immunoreactivity for any of the three neurochemical markers. After analyzing the active firing pattern of the labeled interneurons, we found that in accordance with previous findings PV-positive neurons were always FS interneurons, with either continuous or stuttering spiking. SOM-positive and Prox1-positive interneurons were all non-FS interneurons; these displayed heterogeneous firing patterns with both adapting and non-adapting non-FS behaviors (examples in Fig. 6).

**Figure 6.**
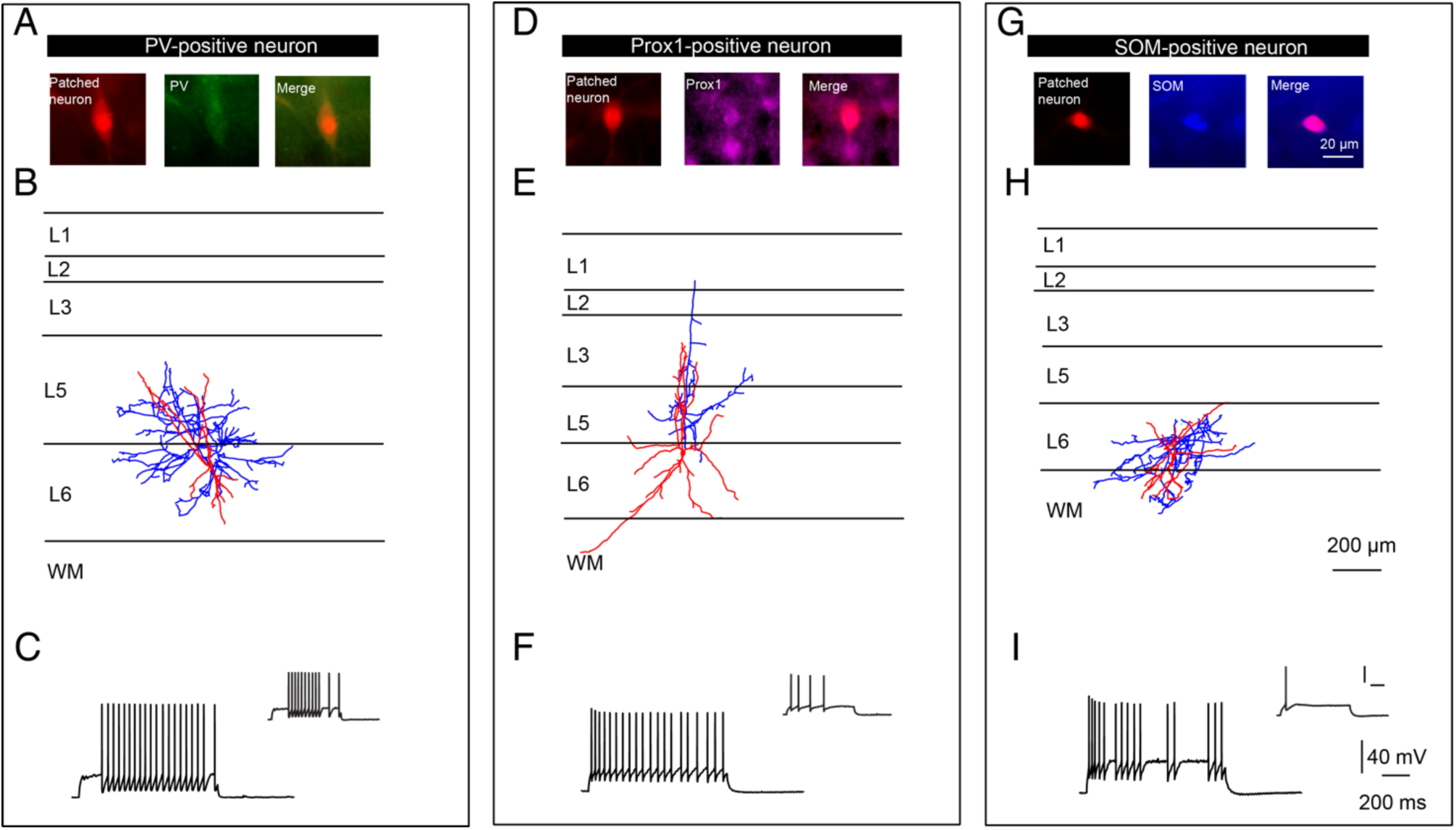
Neurochemical marker expression of individual L6 interneurons in rat mPFC. Whole-cell patch-clamp recordings were made with simultaneous biocytin and Alexa 594 filling (red) to identify the location of patched neurons. Immunostaining was performed after a brief fixation period in PFA to check the expression of PV (green), Prox1 (purple) and SOM (blue) of the patched neurons. Representative examples of interneurons expressing PV (A), Prox1 (D) and somatostatin (G) with morphological reconstructions (B, E, H) and firing pattern (C, F, I, left: the first 10 spikes traces; right: the firing traces at the current 100 pA above the threshold) are shown. Axon is labeled in blue, soma and dendrites in red.

## Discussion

Similar to L6 pyramidal cells in the rat mPFC (Van Aerde and Feldmeyer 2015), L6 interneurons showed a high degree of morphological diversity, ranging from small neurons that were exclusively confined to their home layer to neurons with axons projecting to superficial layers terminating in L1. A CA based on the morphological properties, in particular the axonal projection pattern and soma position revealed 4 clusters, C1 to C4. C1 and C2 displayed translaminar axonal collaterals mainly outside the home layer, while C3 and C4 showed local axonal projections within L6 and WM. C1 and C2 were diverse with respect to their axonal projections, where C1 spanned all cortical layers except L6, while the latter terminated mainly in L5. Similarly, C3 was exclusively restricted to L6, while C4 showed projections also to WM. Interestingly, the axonal projection patterns of L6 interneurons were closely related to their soma position and a correlation between morphological and electrophysiological properties was also detected for some L6 interneuron clusters.

### Classification of interneurons in L6 of other cortices

Using classification methods (CA based on the axonal properties) similar to those used in our work, a study of L6 interneurons in somatosensory barrel cortex identified five independent groups, including interneurons with local or translaminar axonal projection (Arzt et al. 2018). The L6 interneuron types identified in that study showed only a partial overlap with those described here. In particular, the ‘L2/3/4 inhibitors’ and ‘L5b inhibitors’ of that study resembled the L1/2/3-projecting and L5-projecting interneurons in PFC, respectively. The interneuron type coined locally (L6)-projecting interneurons by us includes the L6, L5/6 and L6/5 inhibitors and contained both FS and non-FS interneurons. In contrast to this study, no L6/WM projecting interneuron group was found.

In another study on L6 interneurons in barrel cortex the authors identified four different types of interneurons mainly based on AP firing properties and molecular marker expression (Perrenoud et al. 2013). However, because the axonal domain of these interneurons was not analysed quantitatively, it is difficult to compare the L6 interneuron types of that study directly with those found here. Similar to our findings, the authors found that ∼50% of all L6 interneurons were FS, PV-expressing putative L6 small and large basket cells while the remainder were non-FS, adapting firing interneurons expressing neuropeptide Y, SOM and vasoactive intestinal peptide (VIP)-expressing interneurons, respectively.

In a recent study using a correlated analysis of electrophysiological, molecular/transcriptomic and morphological properties of interneurons in mouse visual cortex, several different distinct neuronal cell types were identified (Gouwens et al. 2019). Using single-cell RNA sequencing, the authors describe three subsets of L6 interneurons most of which have a local axonal domain. They did not describe a group of L1/2/3-projecting L6 interneurons although they show a single example of a SOM-positive L6 interneurons that projects to layers 1 and 2/3. PFC L6 interneurons in C2 resemble to some extent the PV-expressing interneurons described by Gouwens et al. (2019); however, a subset of the PV interneurons of that study is located in L6B and appears to be similar to the PFC FS interneurons in clusters 3 and 4. In a follow-up study three different trasnscriptomic types of PV-expressing interneurons were identified in L6 (Hodge et al. 2019). One group of L6 interneurons in visual cortex termed ‘wide axon, small dendrites’ interneurons showed a high degree of similarity with the majority of the C4 L6/WM- projecting interneurons identified here. In visual cortex these interneurons express the lysosomal-associated membrane protein 5 (Lamp5) and are presumably L6 neurogliaform cells (Hodge et al. 2019).

The classification based on transcriptomic features appeared to be correlated better with electrophysiological properties than with the neuronal morphology, consistent with a study in mouse barrel cortex (Perrenoud et al. 2013).

### Differential innervation of mPFC layers by L6 interneurons

Interneurons in mPFC receive synaptic input predominantly from adjacent, granular frontal cortical regions, but also from virtually all regions of the isocortex, e.g. motor cortex, sensory cortex and associative cortex (Pandya and Yeterian 1990; Petrides and Pandya 1999; Ahrlund-Richter et al. 2019; Sun et al. 2019). In L6, interneurons also receive direct thalamocortical input (Beierlein and Connors 2002). Furthermore, in L6 of visual cortex corticothalamic L6 pyramidal cells have been suggested to innervate L6 interneurons with a higher connection probability than corticocortical cells (West et al. 2006). However, the axonal properties and innervation domains of mPFC L6 interneurons have so far not been described quantitatively.

#### L1/2/3-projecting interneurons

This group of interneurons preferentially targets superficial layers, their axons mostly terminate in layers 1-3, resembling in part SOM-expressing Martinotti-like neurons (Fairén 1984; Kawaguchi and Kubota 1997; Wang et al. 2004); alternatively some may also belong to the rare group of L6 VIP cells which do not appear to have the typical bipolar morphology of VIP cells in L2/3 (Perrenoud et al. 2013; Prönneke et al. 2015).

A distinctive feature of L1/2/3-projecting L6 interneurons is that they only sparsely innervate their home layer because they have only few collaterals in L6; most of their axon collaterals reside in other layers, projecting in a largely vertical fashion. This suggests that L1/2/3- projecting interneurons form only little if any synaptic connections within their home layer, but may establish distal dendritic inhibition in L1 of almost all pyramidal cells from different layers (Silberberg and Markram 2007; Higley 2014; Yavorska and Wehr 2016). In addition, they may also provide inhibition to apical and basal dendrites of both L3 and L5 pyramidal cells. They are therefore in a position to suppress the NMDA spike zone on apical tufts and Ca^2+^ spike zone on distal apical dendrites of pyramidal cells (Larkum et al. 2009; Palmer et al. 2014). L1/2/3- projecting interneurons are the smallest population of our sample of 48 interneurons, a finding similar to that for L6 somatosensory cortex (Arzt et al. 2018).

#### L5-projecting interneurons

As found for L1/2/3-projecting interneurons, this cluster of neurons mainly targets neurons outside the home layer with most of their axonal collaterals located in L5. It is therefore likely that these neurons target the basal dendrites of pyramidal cells in L5 and in particular L5B. They may also innervate the proximal portion of the apical dendrites of L6 upright pyramidal cells. The axonal projection pattern of L5-projecting interneurons were similar with the ‘L5b inhibitors’ found in L6 of somatosensory cortex (Arzt et al. 2018). They may play an important role in determining spike timing during slow oscillation episodes (Van Aerde et al. 2009).

#### Locally-projecting interneurons

These mPFC L6 interneurons are located close to the middle portion of L6 and innervate mainly local neurons in L6. Potential target structures in L6 are basal dendrites of upright pyramidal cells, main and basal dendrites of inverted pyramidal cells as well as other interneurons. Locally-projecting L6 interneurons show both FS and non-FS firing patterns. Based on their morphological features, locally-projecting FS L6 interneurons are reminiscent of FS basket cells while locally-projecting non-FS L6 interneurons could be the radially projecting neurogliaform-like cells (Kawaguchi and Kubota 1993; Wang et al. 2002; Armstrong et al. 2012; Overstreet-Wadiche and Mcbain 2015; Lagler et al. 2016; Gouwens et al. 2019).

#### L6/WM-projecting interneurons

The last group of L6 interneurons are an extension of the locally-projecting L6 interneurons of C3 that inhibit not only L6 pyramidal cells and interneurons but also send collaterals to the WM. These interneurons were found deep in L6 and show profuse axonal branching in their home layer and the WM. Potential targets of L6/WM-projecting interneurons are basal dendrites of all L6B neurons as well as the ‘main’ dendrites of L6B non-pyramidal excitatory neurons, e.g. inverted, ‘tangentially’ oriented and ‘horizontally’ oriented neurons (Marx and Feldmeyer 2013; Gouwens et al. 2019).

WM interstitial cells are considered to be among the oldest neurons of the cerebral cortex; their density declines during development due to differential growth of the WM and an absolute decrease in subplate neuron number (Chun and Shatz 1989). It has been suggested that persistent GABAergic WM interstitial cells receive excitatory and inhibitory input from cortical and subcortical areas (Von Engelhardt et al. 2011). We found that L6/WM-projecting interneurons extend more than a quarter of their axons into the WM so that they are in a position to provide substantial synaptic input to WM interstitial cells.

### Correlation between interneuron types and electrophysiological properties

In mPFC, L6 interneurons show a large diversity in their electrophysiological properties. Discontinuous/stuttering and continuous FS interneurons, neurons with delayed firing onset, adapting neurons and non-adapting non-FS neurons were all identified in this area, in accordance with findings in other brain regions (Kawaguchi 1993; Cauli et al. 1997; Ascoli et al. 2008; Defelipe et al. 2013; Feldmeyer et al. 2018). In our study, we found no correlation between axonal projection and passive membrane properties in agreement with previous studies of L6 in somatosensory cortex (Kumar and Ohana 2008; Arzt et al. 2018). However, a correlation between firing pattern and morphology was found for interneurons of morphological C1 and C2: all L1/2/3-projecting interneurons were adapting neurons and all L5-projecting interneurons were FS interneurons. Interneurons in C3 and C4 showed a heterogeneous firing pattern and the combined CA revealed sub-groups in these two clusters. Interneurons in C3 can be divided into FS and non-FS locally-projecting interneuron (Suppl. Fig. 3D) and those in C4 into FS and non-FS L6/WM-projecting interneurons. Notably, non-FS but not FS L6/WM-projecting interneurons have a single axon branch that extend to L2/3 (Suppl. Fig. 3E) suggesting that electrophysiological parameters may help to reveal the existence of interneuron subtypes.

Perrenoud and coworkers (Perrenoud et al. 2013) have classified barrel cortex L6 interneurons based on electrophysiological and molecular but not morphological properties. When the authors used somatodendritic morphologies as an additional CA parameter this produced clusters that were largely overlapping with those generated using electrophysiological and molecular properties alone. However, a correlation between electrophysiology and axonal morphologies was only found in a study on L2/3 interneurons in frontal cortex (Kawaguchi 1995), suggesting that interneurons with specific axonal projection patterns might have specific AP firing properties.

In contrast, no correlation between the electrophysiological properties of interneurons and their axonal and/or dendritic morphologies was found in several other studies (Gupta et al. 2000; Kumar and Ohana 2008; Helmstaedter et al. 2009; Arzt et al. 2018; Emmenegger et al. 2018). However, electrophysiological properties of a neuron may not be stable even over short time periods because they are under the influence of neuromodulatory transmitters such as acetylcholine, dopamine, serotonin etc. acting via G-protein coupled receptors which affect both neuronal excitability by up- or down-regulation of ion channels such as Kir, KCa, and HCN channels (Dong et al. 2004; Fontanez and Porter 2006; Gulledge and Kawaguchi 2007; Kruglikov and Rudy 2008; Satake et al. 2008; Eggermann and Feldmeyer 2009; Celada et al. 2013; Yi et al. 2013; Van Aerde et al. 2015; Kahnt and Tobler 2017; Qi et al. 2017; Prönneke et al. 2019).

Neurochemical markers are a good indicator of neuronal cell types but recent studies using single cell RNA sequencing showed an ever increasing diversity within the three major groups of PV-. SOM-, and VIP-expressing cortical interneurons (Zeisel et al. 2015; Johnson and Walsh 2017; Paul et al. 2017; Tasic et al. 2018; Gouwens et al. 2019). Here, the expression of the molecular markers PV, SOM and Prox1 in PFC L6 interneurons showed only a weak correlation with the axonal projection pattern. However, there was a correlation between morphology and AP firing pattern for a subset of mPFC L6 interneurons.

Using the axonal projection pattern to identify interneuron types, our analysis provides information about the potential connectivity and density of synaptic contacts of L6 interneurons in mPFC. We found that most L6 interneurons are capable to inhibit target neurons in L5 and L6. However, the L1/2/3-projecting L6 interneurons (10% of the L6 interneuron population)) are in a position to innervate neurons located in superficial layers or apical dendrites of deep layer pyramidal cells. Correlations were found between neuron location and axonal distribution as well as between morphology and intrinsic electrophysiology (partially with C1 and C2), suggesting that interneurons located in the same sub-region of L6 have similar intrinsic properties, so that they may have a similar function in the local microcircuitry. However, the parameters we chose for the morphological classification are certainly not exhaustive and therefore likely to miss some of the properties of L6 interneurons. For example, in morphological C3 and C4, some neurons have a dense while others have only a sparse axon plexus; in addition, a few interneurons showed a horizontal axonal projection pattern. These findings suggest that potential sub-clusters exist within C3 and C4 (see also above). Multiple recordings between L6 interneurons synaptically coupled to excitatory neurons and GABAergic interneurons in different layers may help to determine the specific innervation domains of L6 interneurons and therefore enable us to better understand the role of L6 interneurons in the local microcircuitry.

## Acknowledgement

We thank Werner Hucko for excellent technical assistance and Dr. Karlijn van Aerde for custom-written macros in Igor Pro software. We warmly thank Dr. Guanxiao Qi for helpful discussions.

## Author contributions

D.F., C.D. and V.E. designed the research; C.D., V.E. and K.S. performed experiments and data analysis; C.D., V.E. and D.F. wrote the manuscript. The authors declare no conflicts of interest.

## References

Ahrlund-Richter S, Xuan Y, van Lunteren JA, Kim H, Ortiz C, Pollak Dorocic I, Meletis K, Carlen M. 2019. A whole-brain atlas of monosynaptic input targeting four different cell types in the medial prefrontal cortex of the mouse. Nat Neurosci 22:657–668.

Armstrong C, Krook-Magnuson E, Soltesz I. 2012. Neurogliaform and Ivy Cells: A Major Family of nNOS Expressing GABAergic Neurons. Front Neural Circuits 6:23.

Arzt M, Sakmann B, Meyer HS. 2018. Anatomical Correlates of Local, Translaminar, and Transcolumnar Inhibition by Layer 6 GABAergic Interneurons in Somatosensory Cortex. Cereb Cortex 28:2763–2774.

Ascoli GA, Alonso-Nanclares L, Anderson SA, Barrionuevo G, Benavides-Piccione R, Burkhalter A, Buzsaki G, Cauli B, Defelipe J, Fairen A, Feldmeyer D, Fishell G, Fregnac Y, Freund TF, Gardner D, Gardner EP, Goldberg JH, Helmstaedter M, Hestrin S, Karube F, Kisvarday ZF, Lambolez B, Lewis DA, Marin O, Markram H, Munoz A, Packer A, Petersen CC, Rockland KS, Rossier J, Rudy B, Somogyi P, Staiger JF, Tamas G, Thomson AM, Toledo-Rodriguez M, Wang Y, West DC, Yuste R. 2008. Petilla terminology: nomenclature of features of GABAergic interneurons of the cerebral cortex. Nature reviews Neuroscience 9:557–568.

Beierlein M, Connors BW. 2002. Short-term dynamics of thalamocortical and intracortical synapses onto layer 6 neurons in neocortex. J Neurophysiol 88:1924–1932.

Benes FM, Berretta S. 2001. GABAergic interneurons: Implications for understanding schizophrenia and bipolar disorder (vol 25, pg 1, 2001). Neuropsychopharmacology 25:453–453.

Beneyto M, Abbott A, Hashimoto T, Lewis DA. 2011. Lamina-specific alterations in cortical GABA(A) receptor subunit expression in schizophrenia. Cereb Cortex 21:999–1011.

Cauli B, Audinat E, Lambolez B, Angulo MC, Ropert N, Tsuzuki K, Hestrin S, Rossier J. 1997. Molecular and physiological diversity of cortical nonpyramidal cells. J Neurosci 17:3894–3906.

Celada P, Puig MV, Artigas F. 2013. Serotonin modulation of cortical neurons and networks. Front Integr Neurosci 7:25.

Chun JJ, Shatz CJ. 1989. Interstitial cells of the adult neocortical white matter are the remnant of the early generated subplate neuron population. J Comp Neurol 282:555–569.

DeFelipe J, Lopez-Cruz PL, Benavides-Piccione R, Bielza C, Larranaga P, Anderson S, Burkhalter A, Cauli B, Fairen A, Feldmeyer D, Fishell G, Fitzpatrick D, Freund TF, Gonzalez-Burgos G, Hestrin S, Hill S, Hof PR, Huang J, Jones EG, Kawaguchi Y, Kisvarday Z, Kubota Y, Lewis DA, Marin O, Markram H, McBain CJ, Meyer HS, Monyer H, Nelson SB, Rockland K, Rossier J, Rubenstein JL, Rudy B, Scanziani M, Shepherd GM, Sherwood CC, Staiger JF, Tamas G, Thomson A, Wang Y, Yuste R, Ascoli GA. 2013. New insights into the classification and nomenclature of cortical GABAergic interneurons. Nature reviews Neuroscience 14:202–216.

Dong Y, Cooper D, Nasif F, Hu XT, White FJ. 2004. Dopamine modulates inwardly rectifying potassium currents in medial prefrontal cortex pyramidal neurons. J Neurosci 24:3077–3085.

Eggermann E, Feldmeyer D. 2009. Cholinergic filtering in the recurrent excitatory microcircuit of cortical layer 4. Proc Natl Acad Sci U S A 106:11753–11758.

Elston GN. 2003. Cortex, cognition and the cell: new insights into the pyramidal neuron and prefrontal function. Cerebral cortex 13:1124–1138.

Emmenegger V, Qi G, Wang H, Feldmeyer D. 2018. Morphological and Functional Characterization of Non-fast-Spiking GABAergic Interneurons in Layer 4 Microcircuitry of Rat Barrel Cortex. Cereb Cortex 28:1439–1457.

Euston DR, Gruber AJ, McNaughton BL. 2012. The role of medial prefrontal cortex in memory and decision making. Neuron 76:1057–1070.

Fairén A, DeFelipe, J., and Regidor, J. 1984. Nonpyramidal Neurons. In Cerebral Cortex: Cellular Components of the Cerebral Cortex:206–211.

Feldmeyer D, Qi G, Emmenegger V, Staiger JF. 2018. Inhibitory interneurons and their circuit motifs in the many layers of the barrel cortex. Neuroscience 368:132–151.

Ferguson BR, Gao WJ. 2018. PV Interneurons: Critical Regulators of E/I Balance for Prefrontal Cortex-Dependent Behavior and Psychiatric Disorders. Front Neural Circuits 12:37.

Fontanez DE, Porter JT. 2006. Adenosine A1 receptors decrease thalamic excitation of inhibitory and excitatory neurons in the barrel cortex. Neuroscience 137:1177–1184.

Fuster JM. 2001. The prefrontal cortex--an update: time is of the essence. Neuron 30:319–333.

Gabbott PL, Dickie BG, Vaid RR, Headlam AJ, Bacon SJ. 1997. Local-circuit neurones in the medial prefrontal cortex (areas 25, 32 and 24b) in the rat: morphology and quantitative distribution. J Comp Neurol 377:465–499.

Gabbott PL, Warner TA, Jays PR, Salway P, Busby SJ. 2005. Prefrontal cortex in the rat: projections to subcortical autonomic, motor, and limbic centers. J Comp Neurol 492:145–177.

Gouwens NW, Sorensen SA, Berg J, Lee C, Jarsky T, Ting J, Sunkin SM, Feng D, Anastassiou CA, Barkan E, Bickley K, Blesie N, Braun T, Brouner K, Budzillo A, Caldejon S, Casper T, Castelli D, Chong P, Crichton K, Cuhaciyan C, Daigle TL, Dalley R, Dee N, Desta T, Ding SL, Dingman S, Doperalski A, Dotson N, Egdorf T, Fisher M, de Frates RA, Garren E, Garwood M, Gary A, Gaudreault N, Godfrey K, Gorham M, Gu H, Habel C, Hadley K, Harrington J, Harris JA, Henry A, Hill D, Josephsen S, Kebede S, Kim L, Kroll M, Lee B, Lemon T, Link KE, Liu X, Long B, Mann R, McGraw M, Mihalas S, Mukora A, Murphy GJ, Ng L, Ngo K, Nguyen TN, Nicovich PR, Oldre A, Park D, Parry S, Perkins J, Potekhina L, Reid D, Robertson M, Sandman D, Schroedter M, Slaughterbeck C, Soler-Llavina G, Sulc J, Szafer A, Tasic B, Taskin N, Teeter C, Thatra N, Tung H, Wakeman W, Williams G, Young R, Zhou Z, Farrell C, Peng H, Hawrylycz MJ, Lein E, Ng L, Arkhipov A, Bernard A, Phillips JW, Zeng H, Koch C. 2019. Classification of electrophysiological and morphological neuron types in the mouse visual cortex. Nat Neurosci 22:1182–1195.

Guillery RW, Sherman SM. 2002. Thalamic relay functions and their role in corticocortical communication: generalizations from the visual system. Neuron 33:163–175.

Gulledge AT, Kawaguchi Y. 2007. Phasic cholinergic signaling in the hippocampus: functional homology with the neocortex? Hippocampus 17:327–332.

Gupta A, Wang Y, Markram H. 2000. Organizing principles for a diversity of GABAergic interneurons and synapses in the neocortex. Science 287:273–278.

Helmstaedter M, Sakmann B, Feldmeyer D. 2009. The relation between dendritic geometry, electrical excitability, and axonal projections of L2/3 interneurons in rat barrel cortex. Cereb Cortex 19:938–950.

Higley MJ. 2014. Localized GABAergic inhibition of dendritic Ca(2+) signalling. Nat Rev Neurosci 15:567–572.

Hirai Y, Morishima M, Karube F, Kawaguchi Y. 2012. Specialized cortical subnetworks differentially connect frontal cortex to parahippocampal areas. J Neurosci 32:1898–1913.

Hodge RD, Bakken TE, Miller JA, Smith KA, Barkan ER, Graybuck LT, Close JL, Long B, Johansen N, Penn O, Yao ZZ, Eggermont J, Hollt T, Levi BP, Shehata SI, Aevermann B, Beller A, Bertagnolli D, Brouner K, Casper T, Cobbs C, Dalley R, Dee N, Ding SL, Ellenbogen RG, Fong O, Garren E, Goldy J, Gwinn RP, Hirschstein D, Keene CD, Keshk M, Ko AL, Lathia K, Mahfouz A, Maltzer Z, McGraw M, Nguyen TN, Nyhus J, Ojemann JG, Oldre A, Parry S, Reynolds S, Rimorin C, Shapovalova NV, Somasundaram S, Szafer A, Thomsen ER, Tieu M, Quon G, Scheuermann RH, Yuste R, Sunkin SM, Lelieveldt B, Feng D, Ng L, Bernard A, Hawrylycz M, Phillips JW, Tasic B, Zeng HK, Jones AR, Koch C, Lein ES. 2019. Conserved cell types with divergent features in human versus mouse cortex. Nature 573:61–68.

Johnson MB, Walsh CA. 2017. Cerebral cortical neuron diversity and development at single-cell resolution. Curr Opin Neurobiol 42:9–16.

Kahnt T, Tobler PN. 2017. Dopamine Modulates the Functional Organization of the Orbitofrontal Cortex. The Journal of neuroscience : the official journal of the Society for Neuroscience 37:1493–1504.

Kawaguchi Y. 1993. Physiological, morphological, and histochemical characterization of three classes of interneurons in rat neostriatum. J Neurosci 13:4908–4923.

Kawaguchi Y. 1995. Physiological subgroups of nonpyramidal cells with specific morphological characteristics in layer II/III of rat frontal cortex. J Neurosci 15:2638–2655.

Kawaguchi Y, Kubota Y. 1993. Correlation of physiological subgroupings of nonpyramidal cells with parvalbumin- and calbindinD28k-immunoreactive neurons in layer V of rat frontal cortex. Journal of neurophysiology 70:387–396.

Kawaguchi Y, Kubota Y. 1997. GABAergic cell subtypes and their synaptic connections in rat frontal cortex. Cerebral cortex 7:476–486.

Kepecs A, Fishell G. 2014. Interneuron cell types are fit to function. Nature 505:318–326.

Kruglikov I, Rudy B. 2008. Perisomatic GABA release and thalamocortical integration onto neocortical excitatory cells are regulated by neuromodulators. Neuron 58:911–924.

Kubota Y, Hatada S, Kondo S, Karube F, Kawaguchi Y. 2007. Neocortical inhibitory terminals innervate dendritic spines targeted by thalamocortical afferents. J Neurosci 27:1139–1150.

Kumar P, Ohana O. 2008. Inter- and intralaminar subcircuits of excitatory and inhibitory neurons in layer 6a of the rat barrel cortex. J Neurophysiol 100:1909–1922.

Lagler M, Ozdemir AT, Lagoun S, Malagon-Vina H, Borhegyi Z, Hauer R, Jelem A, Klausberger T. 2016. Divisions of Identified Parvalbumin-Expressing Basket Cells during Working Memory-Guided Decision Making. Neuron 91:1390–1401.

Larkum ME, Nevian T, Sandler M, Polsky A, Schiller J. 2009. Synaptic integration in tuft dendrites of layer 5 pyramidal neurons: a new unifying principle. Science 325:756–760.

Lein ES, Belgard TG, Hawrylycz M, Molnar Z. 2017. Transcriptomic Perspectives on Neocortical Structure, Development, Evolution, and Disease. Annual Review of Neuroscience, Vol 40 40:629–652.

Lübke J, Roth A, Feldmeyer D, Sakmann B. 2003. Morphometric analysis of the columnar innervation domain of neurons connecting layer 4 and layer 2/3 of juvenile rat barrel cortex. Cereb Cortex 13:1051–1063.

Markram H, Toledo-Rodriguez M, Wang Y, Gupta A, Silberberg G, Wu C. 2004. Interneurons of the neocortical inhibitory system. Nat Rev Neurosci 5:793–807.

Marx M, Feldmeyer D. 2013. Morphology and physiology of excitatory neurons in layer 6b of the somatosensory rat barrel cortex. Cerebral cortex 23:2803–2817.

Marx M, Günter RH, Hucko W, Radnikow G, Feldmeyer D. 2012. Improved biocytin labeling and neuronal 3D reconstruction. Nat Protoc 7:394–407.

Miller EK, Cohen JD. 2001. An integrative theory of prefrontal cortex function. Annual review of neuroscience 24:167–202.

Möhler H. 2012. The GABA system in anxiety and depression and its therapeutic potential. Neuropharmacology 62:42–53.

Overstreet-Wadiche L, McBain CJ. 2015. Neurogliaform cells in cortical circuits. Nat Rev Neurosci 16:458–468.

Palmer L, Murayama M, Larkum M. 2012. Inhibitory regulation of dendritic activity in vivo. Front Neural Circuits 6:26.

Palmer LM, Shai AS, Reeve JE, Anderson HL, Paulsen O, Larkum ME. 2014. NMDA spikes enhance action potential generation during sensory input. Nature Neuroscience 17:383.

Pandya DN, Yeterian EH. 1990. Prefrontal cortex in relation to other cortical areas in rhesus monkey: architecture and connections. Progress in brain research 85:63–94.

Paul A, Crow M, Raudales R, He M, Gillis J, Huang ZJ. 2017. Transcriptional Architecture of Synaptic Communication Delineates GABAergic Neuron Identity. Cell 171:522–539.e520.

Perrenoud Q, Rossier J, Geoffroy H, Vitalis T, Gallopin T. 2013. Diversity of GABAergic interneurons in layer VIa and VIb of mouse barrel cortex. Cereb Cortex 23:423–441.

Petrides M, Pandya DN. 1999. Dorsolateral prefrontal cortex: comparative cytoarchitectonic analysis in the human and the macaque brain and corticocortical connection patterns. The European journal of neuroscience 11:1011–1036.

Prönneke A, Scheuer B, Wagener RJ, Mock M, Witte M, Staiger JF. 2015. Characterizing VIP Neurons in the Barrel Cortex of VIPcre/tdTomato Mice Reveals Layer-Specific Differences. Cereb Cortex 25:4854–4868.

Prönneke A, Witte M, Möck M, Staiger JF. 2019. Neuromodulation Leads to a Burst-Tonic Switch in a Subset of VIP Neurons in Mouse Primary Somatosensory (Barrel) Cortex. Cereb Cortex.

Qi G, van Aerde K, Abel T, Feldmeyer D. 2017. Adenosine Differentially Modulates Synaptic Transmission of Excitatory and Inhibitory Microcircuits in Layer 4 of Rat Barrel Cortex. Cereb Cortex 27:4411–4422.

Riga D, Matos MR, Glas A, Smit AB, Spijker S, Van den Oever MC. 2014. Optogenetic dissection of medial prefrontal cortex circuitry. Front Syst Neurosci 8:230.

Rudy B, Fishell G, Lee S, Hjerling-Leffler J. 2011. Three groups of interneurons account for nearly 100% of neocortical GABAergic neurons. Dev Neurobiol 71:45–61.

Satake T, Mitani H, Nakagome K, Kaneko K. 2008. Individual and additive effects of neuromodulators on the slow components of afterhyperpolarization currents in layer V pyramidal cells of the rat medial prefrontal cortex. Brain Res 1229:47–60.

Silberberg G, Markram H. 2007. Disynaptic inhibition between neocortical pyramidal cells mediated by Martinotti cells. Neuron 53:735–746.

Sun Q, Li X, Ren M, Zhao M, Zhong Q, Ren Y, Luo P, Ni H, Zhang X, Zhang C, Yuan J, Li A, Luo M, Gong H, Luo Q. 2019. A whole-brain map of long-range inputs to GABAergic interneurons in the mouse medial prefrontal cortex. Nat Neurosci 22:1357–1370.

Tasic B, Yao Z, Graybuck LT, Smith KA, Nguyen TN, Bertagnolli D, Goldy J, Garren E, Economo MN, Viswanathan S, Penn O, Bakken T, Menon V, Miller J, Fong O, Hirokawa KE, Lathia K, Rimorin C, Tieu M, Larsen R, Casper T, Barkan E, Kroll M, Parry S, Shapovalova NV, Hirschstein D, Pendergraft J, Sullivan HA, Kim TK, Szafer A, Dee N, Groblewski P, Wickersham I, Cetin A, Harris JA, Levi BP, Sunkin SM, Madisen L, Daigle TL, Looger L, Bernard A, Phillips J, Lein E, Hawrylycz M, Svoboda K, Jones AR, Koch C, Zeng H. 2018. Shared and distinct transcriptomic cell types across neocortical areas. Nature 563:72–78.

Thomson AM. 2010. Neocortical layer 6, a review. Front Neuroanat 4:13.

Thomson AM, West DC, Wang Y, Bannister AP. 2002. Synaptic connections and small circuits involving excitatory and inhibitory neurons in layers 2-5 of adult rat and cat neocortex: triple intracellular recordings and biocytin labelling in vitro. Cereb Cortex 12:936–953.

Tremblay R, Lee S, Rudy B. 2016. GABAergic Interneurons in the Neocortex: From Cellular Properties to Circuits. Neuron 91:260–292.

Uylings HB, Groenewegen HJ, Kolb B. 2003a. Do rats have a prefrontal cortex? Behav Brain Res 146:3–17.

Uylings HB, Groenewegen HJ, Kolb B. 2003b. Do rats have a prefrontal cortex? Behav Brain Research 146:3–17.

Uylings HB, van Eden CG. 1990. Qualitative and quantitative comparison of the prefrontal cortex in rat and in primates, including humans. Prog Brain Res 85:31–62.

van Aerde KI, Feldmeyer D. 2015. Morphological and physiological characterization of pyramidal neuron subtypes in rat medial prefrontal cortex. Cereb Cortex 25:788–805.

van Aerde KI, Mann EO, Canto CB, Heistek TS, Linkenkaer-Hansen K, Mulder AB, van der Roest M, Paulsen O, Brussaard AB, Mansvelder HD. 2009. Flexible spike timing of layer 5 neurons during dynamic beta oscillation shifts in rat prefrontal cortex. J Physiol 587:5177–5196.

van Aerde KI, Qi G, Feldmeyer D. 2015. Cell type-specific effects of adenosine on cortical neurons. Cerebral cortex 25:772–787.

Van Eden CG, Uylings HB. 1985. Cytoarchitectonic development of the prefrontal cortex in the rat. J Comp Neurol 241:253–267.

Volk DW, Lewis DA. 2005. GABA Targets for the Treatment of Cognitive Dysfunction in Schizophrenia. Curr Neuropharmacol 3:45–62.

von Engelhardt J, Khrulev S, Eliava M, Wahlster S, Monyer H. 2011. 5-HT(3A) receptor-bearing white matter interstitial GABAergic interneurons are functionally integrated into cortical and subcortical networks. J Neurosci 31:16844–16854.

Wang Y, Gupta A, Toledo-Rodriguez M, Wu CZ, Markram H. 2002. Anatomical, physiological, molecular and circuit properties of nest basket cells in the developing somatosensory cortex. Cereb Cortex 12:395–410.

Wang Y, Toledo-Rodriguez M, Gupta A, Wu CZ, Silberberg G, Luo JY, Markram H. 2004. Anatomical, physiological and molecular properties of Martinotti cells in the somatosensory cortex of the juvenile rat. J Physiol-London 561:65–90.

Ward JH. 1963. Hierarchical grouping to optimize an objective function. J Am Stat Assoc 58:236–244.

West DC, Mercer A, Kirchhecker S, Morris OT, Thomson AM. 2006. Layer 6 cortico-thalamic pyramidal cells preferentially innervate interneurons and generate facilitating EPSPs. Cereb Cortex 16:200–211.

Wimmer RD, Schmitt LI, Davidson TJ, Nakajima M, Deisseroth K, Halassa MM. 2015. Thalamic control of sensory selection in divided attention. Nature 526:705–709.

Yang CR, Seamans JK, Gorelova N. 1996. Electrophysiological and morphological properties of layers V-VI principal pyramidal cells in rat prefrontal cortex in vitro. The Journal of neuroscience : the official journal of the Society for Neuroscience 16:1904–1921.

Yavorska I, Wehr M. 2016. Somatostatin-Expressing Inhibitory Interneurons in Cortical Circuits. Front Neural Circuits 10:76.

Yi F, Zhang XH, Yang CR, Li BM. 2013. Contribution of dopamine d1/5 receptor modulation of post-spike/burst afterhyperpolarization to enhance neuronal excitability of layer v pyramidal neurons in prepubertal rat prefrontal cortex. PLoS One 8:e71880.

Yuste R, Hawrylycz M, Aalling N, Arendt D, Armananzas R, Ascoli G, Bielza C, Bokharaie V, Bergmann T, Bystron I, Capogna M, Chang Y, Clemens A, Kock Cd, DeFelipe J, Santos Sd, Dunville K, Feldmeyer D, Fiath R, Fishell G, Foggetti A, Gao X, Ghaderi P, Güntürkun O, Hall VJ, Helmstaedter M, Herculano-Houzel S, Hilscher M, Hirase H, Hjerling-Leffler J, Hodge R, Huang ZJ, Huda R, Juan Y, Khodosevich K, Kiehn O, Koch H, Kuebler E, Kuhnemund M, Larranaga P, Lelieveldt B, Louth EL, Lui J, Mansvelder H, Marín O, Martínez-Trujillo J, Moradi H, Goriounova N, Mohapatra A, Nedergaard M, Němec P, Ofer N, Pfisterer U, Pontes S, Redmond W, Rossier J, Sanes J, Scheuermann R, Saiz ES, Somogyi P, Tamás G, Tolias A, Tosches M, Garcia MT, Aguilar-Valles A, Munguba H, Wozny C, Wuttke T, Yong L, Zeng H, Lein ES. 2019. A community-based transcriptomics classification and nomenclature of neocortical cell types. arXiv 1909.03083.

Zeisel A, Munoz-Manchado AB, Codeluppi S, Lonnerberg P, La Manno G, Jureus A, Marques S, Munguba H, He L, Betsholtz C, Rolny C, Castelo-Branco G, Hjerling-Leffler J, Linnarsson S. 2015. Brain structure. Cell types in the mouse cortex and hippocampus revealed by single-cell RNA-seq. Science 347:1138–1142.

Zeng H, Sanes JR. 2017. Neuronal cell-type classification: challenges, opportunities and the path forward. Nat Rev Neurosci 18:530–546.

Zikopoulos B, Barbas H. 2006. Prefrontal projections to the thalamic reticular nucleus form a unique circuit for attentional mechanisms. J Neurosci 26:7348–7361.

