## Supplemental Materials (4 Figures) for "Layer-specific inhibitory microcircuits of layer 6 interneurons in rat prefrontal cortex"

### Supplementary Materials

Containing 4 figures with legends

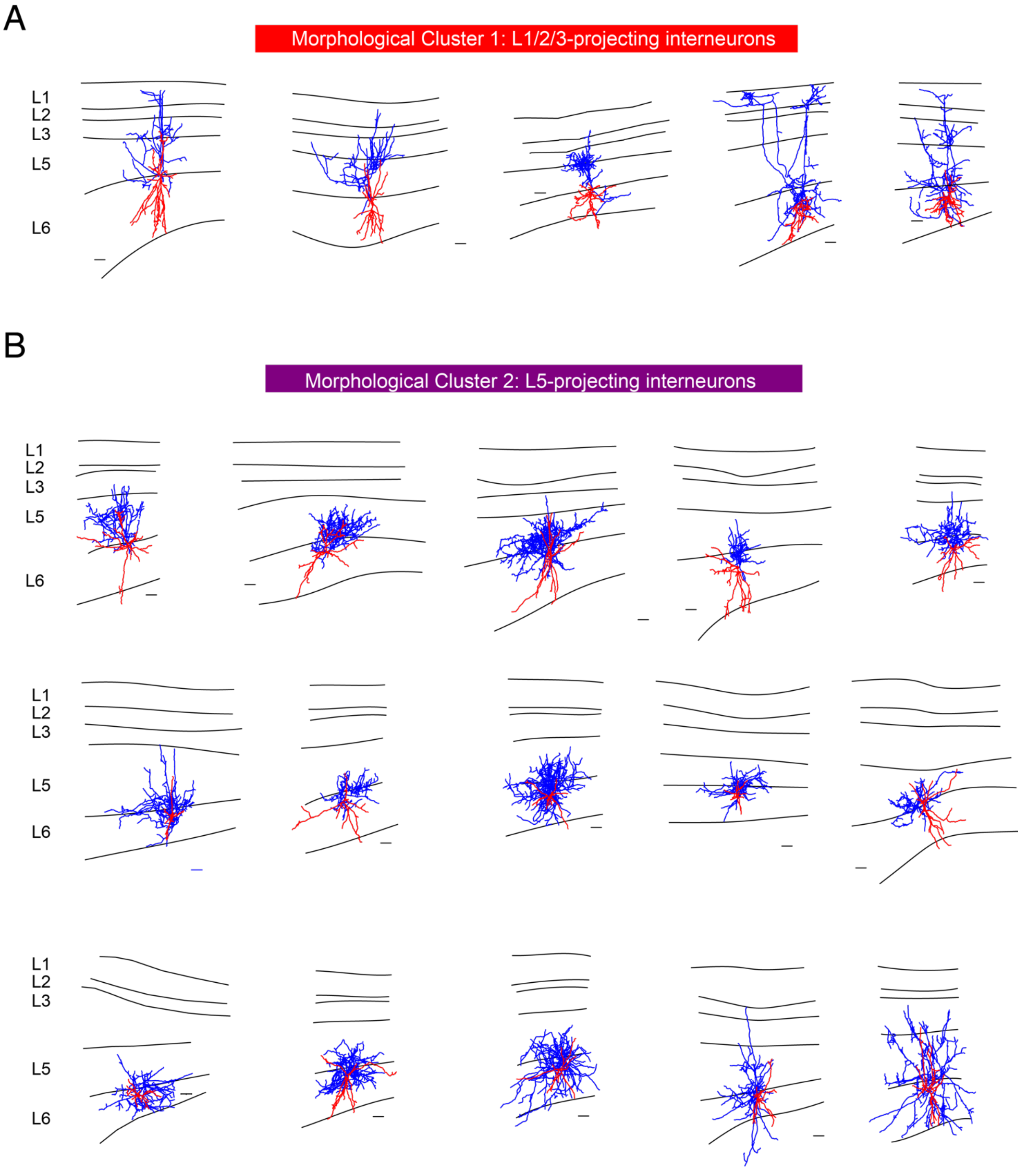


**Supplemental Fig. 1. Morphological cluster 1 and cluster 2 of L6 interneurons.** Individual reconstructions of (A) morphological cluster 1 (L1/2/3-projecting interneurons, n=5) and (B) morphological cluster 2 (L5-projecting interneurons, n=15) revealed by cluster analysis. Dendrites are shown in red and axons in blue. Scale bar, 100 µm.

A

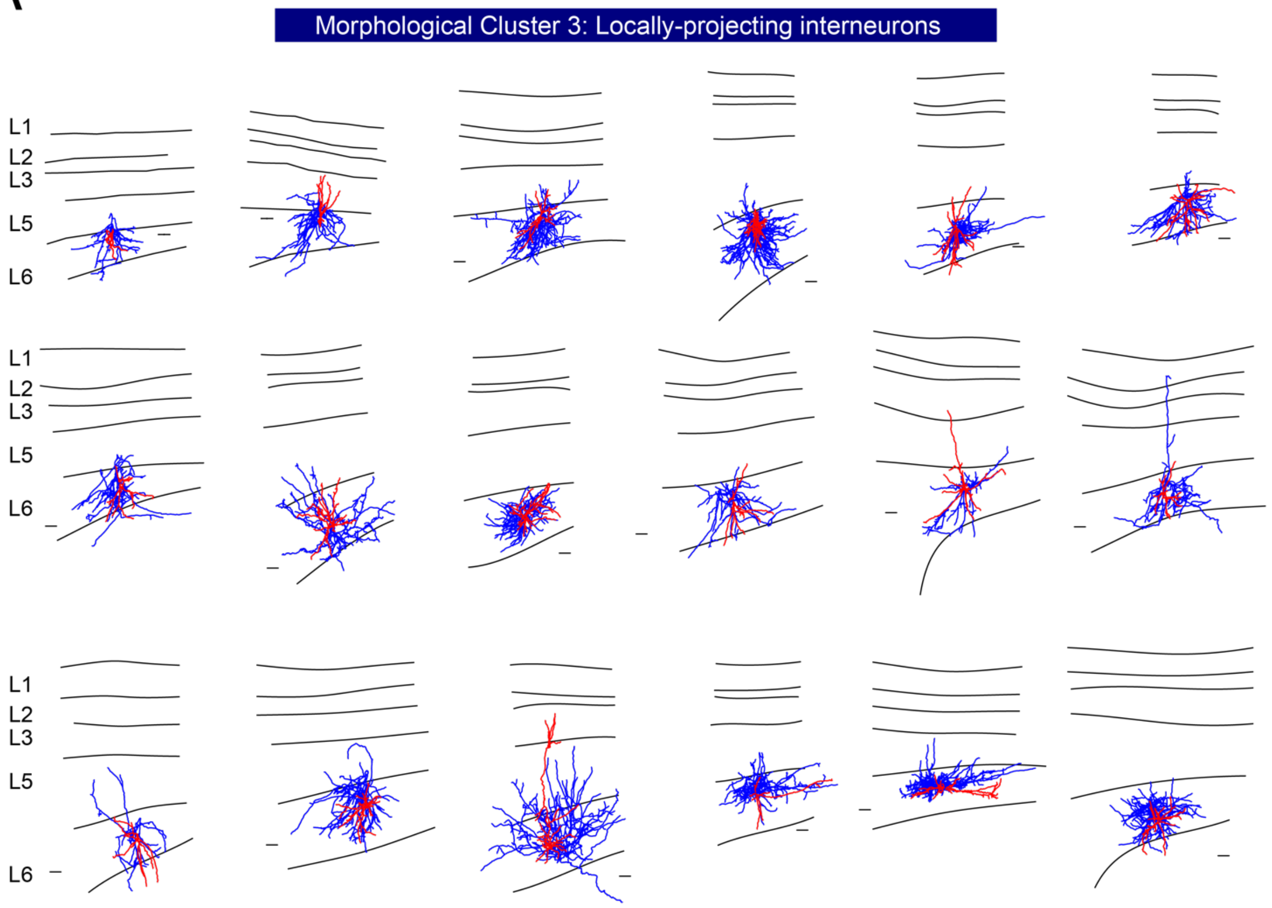
Morphological Cluster 3: Locally-projecting interneurons
L1
L2
L3
L5
L6

B

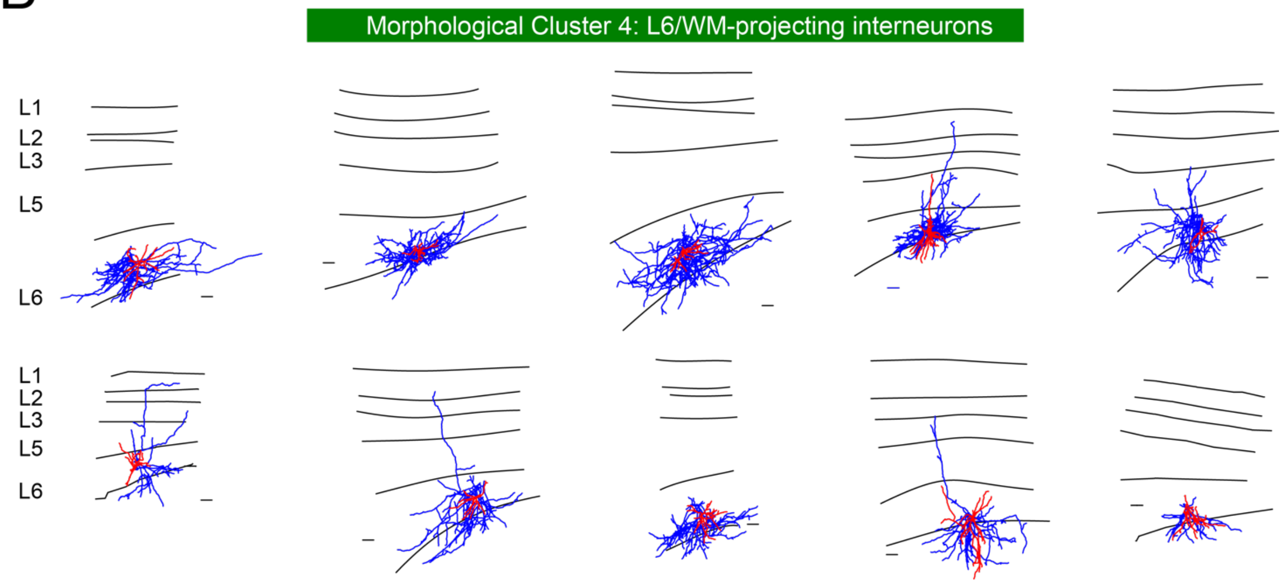
Morphological Cluster 4: L6/WM-projecting interneurons
L1
L2
L3
L5
L6

27

**Supplemental Fig. 2. Morphological cluster 3 and cluster 4 of L6 interneurons.** Individual reconstructions of the (A) morphological cluster 3 (Locally-projecting interneurons, n=18) and (B) morphological cluster 4 (L6/WM-projecting interneurons, n=10) revealed by cluster analysis. Dendrites are shown in red and axons in blue. Scale bar, 100 μm.

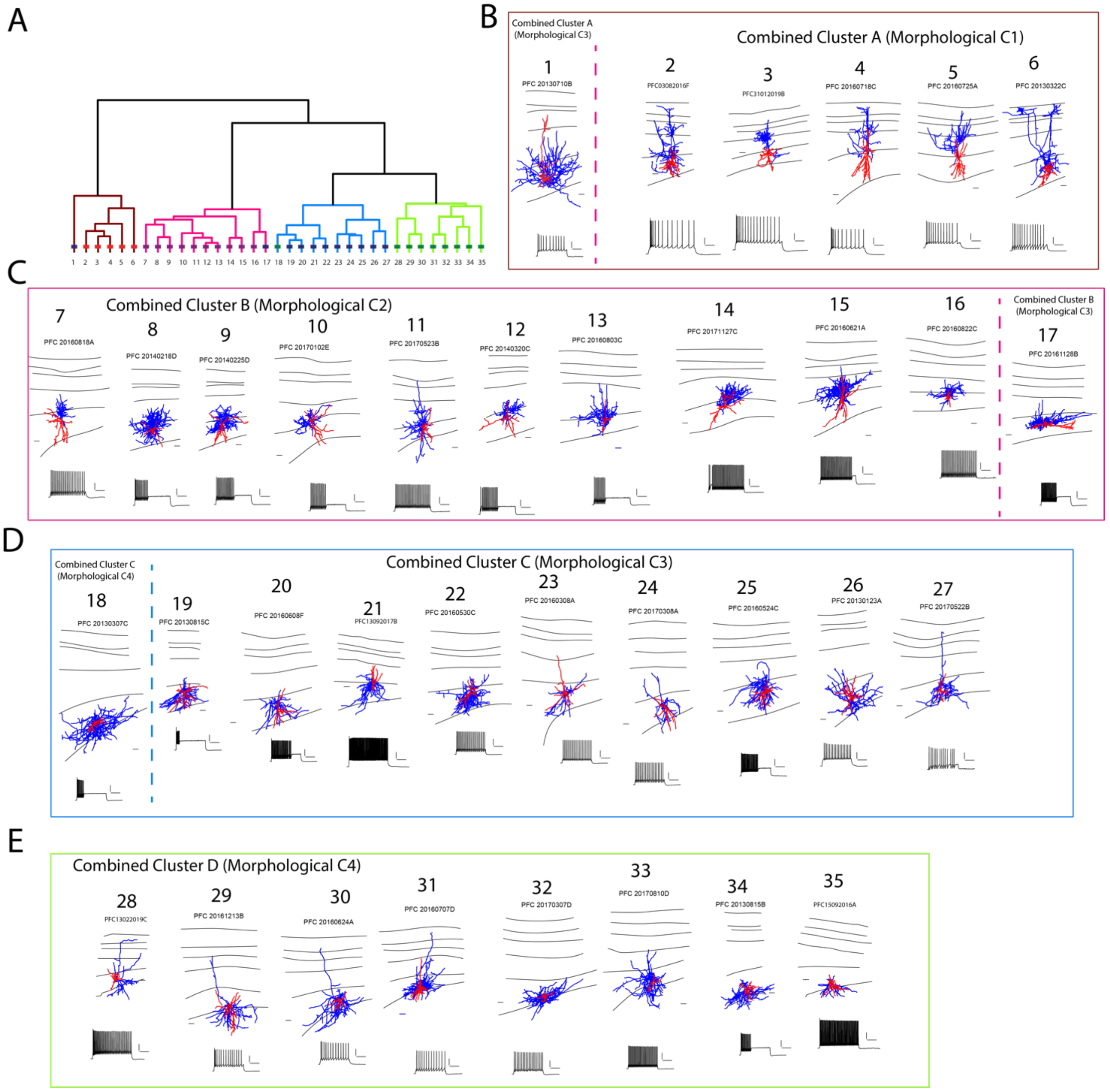


**Supplemental Fig. 3. Morphologies and firing patterns of L6 interneurons in the combined morphological-electrophysiological CA. (A)** Combined CA (details see Fig. 5A) was made by including morphological and membrane active properties. (B-E): Individual morphology and firing pattern of interneurons in each combined cluster. Scale bar, 100 µm for the morphologies, 20 mV and 200 ms for the firing patterns. Potential sub-clusters in cluster C and D were marked in blue/light blue and green/light green respectively.

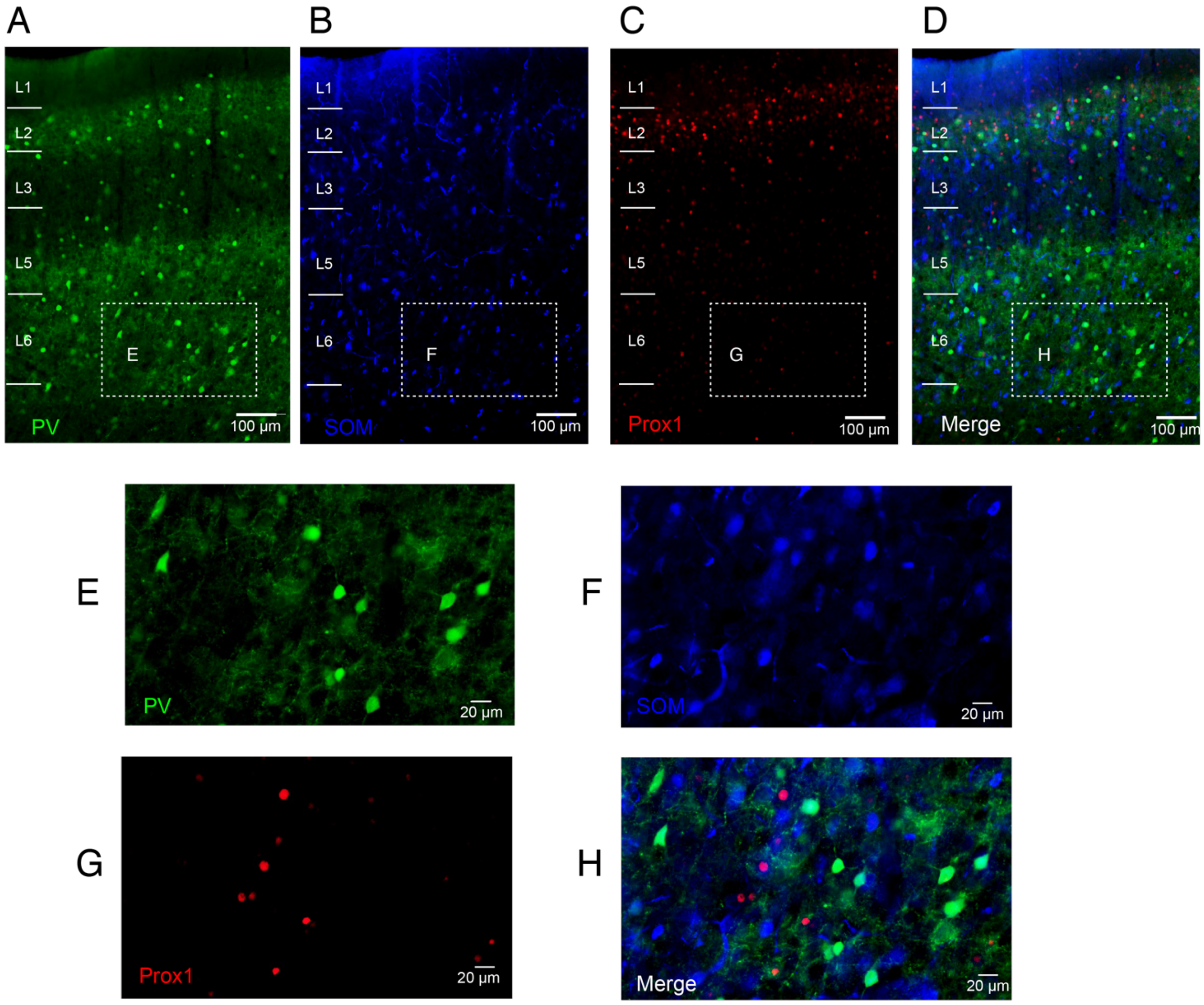


**Supplemental Fig. 4. Expression of neurochemical markers in the same mPFC slice.** Antibody labelling for (A) PV (green), (B) SOM (blue) and (C) Prox1 (red), merged image is showed in D. The area in the dashed box from A-D is enlarged in E-H.
